# Connectomic analysis of thalamus-driven disinhibition in cortical layer 4

**DOI:** 10.1101/2022.06.01.494290

**Authors:** Yunfeng Hua, Sahil Loomba, Verena Pawlak, Philip Laserstein, Kevin M. Boergens, Jason N. D. Kerr, Moritz Helmstaedter

## Abstract

In mammals, sensory signals are transmitted via the thalamus primarily to layer 4 of the primary sensory cortices. While information about average neuronal connectivity in this layer is available, the detailed and higher-order circuit structure is not known. Here, we used 3-dimensional electron microscopy for a connectomic analysis of the thalamus-driven inhibitory network in a layer 4 barrel. We find that thalamic input drives a subset of interneurons with high specificity. These interneurons in turn target spiny stellate and star pyramidal excitatory neurons with subtype specificity. In addition, they create a directed disinhibitory network directly driven by the thalamic input. Together, this circuit can create differential windows of opportunity for activation of the types of excitatory neurons in dependence of strength and timing of thalamic input. With this, we have identified a so-far unknown degree of specialization of the microcircuitry in the main thalamocortical recipient layer of the primary sensory cortex.

## INTRODUCTION

The logic of signal transformation in the main thalamocortical recipient layer of the mammalian brain, layer 4 (L4) of the sensory cortices, has been a major focus of functional (Brecht and Sakmann, 2002; de Kock et al., 2007; Gutnisky et al., 2017; Hubel and Wiesel, 1962; Kanold et al., 2003; Kara et al., 2002; Martinez et al., 2005; Stratford et al., 1996; Yu et al., 2016; Yu et al., 2019) and structural (Ahmed et al., 1994; Ahmed et al., 1997; Egger et al., 2008; Elston et al., 1997; Lübke et al., 2000; Lübke et al., 2003; Motta et al., 2019; Schubert et al., 2003; Staiger et al., 2004; Woolsey and Van der Loos, 1970) investigations. While functional studies have shown the relevance of feedforward inhibition (Bruno and Simons, 2002; Cruikshank et al., 2007; Gabernet et al., 2005; House et al., 2011; Pouille and Scanziani, 2001; Staiger et al., 2009), and recent recordings have demonstrated differential activation of subtypes of interneurons (Yu et al., 2016; Yu et al., 2019), a detailed circuit-level picture of the transformation of the thalamocortical (TC) signal onto excitatory neurons in the cortex is still lacking.

Here, we undertook a connectomic reconstruction of about ¼ of the C2 barrel across all of L4 and parts of L3 and L5A in mouse S1 cortex using serial block-face electron microscopy (SBEM (Denk and Horstmann, 2004)). The tissue had also been imaged using *in vivo* 2-photon laser-scanning microscopy (Denk et al., 1990) prior to the structural experiment. We report findings about the inhibitory synaptic circuitry in L4 that is directly driven by thalamic input, unveiling a high-precision disinhibitory circuitry and a circuit-level distinction between the excitatory neuronal subtypes within L4.

## RESULTS

We used SBEM (Denk and Horstmann, 2004) in continuous imaging mode (Schmidt et al., 2017) to acquire a 3D EM dataset of size 453.2 x 430.4 x 360 µm^3^ at a voxel size of 11.24 x 11.24 x 30 nm^3^, aligned to the C2 barrel (identified by functional 2- photon imaging in-vivo) in an 80-day old mouse (Fig. 1A, tissue staining using(Hua et al., 2015)). The dataset extended from the lower part of L3 to the upper part of L5A (Fig. 1D), containing a total of about 7,600 neuronal cell bodies. We then used our online annotation tool webKnossos (Boergens et al., 2017) to reconstruct the upward pointing (apical) dendrites of n=1,976 neurons in a fraction of the dataset with optimal staining and image alignment (corresponding to about 1/4^th^ of a barrel). The two primary cell types within L4, spiny stellates (SpS) and star pyramidals (StP) (Feldmeyer et al., 1999; LeVay, 1973; Lorente De No, 1938; Lübke et al., 2000; McCormick et al., 1985; Simons and Woolsey, 1984b; White, 1978; Woolsey et al., 1975) were distinguishable based on the existence or lack of an apical dendrite (Fig. 1E). The two excitatory cell types showed surprisingly clear depth separation within L4: SpS occupied the lower two third of L4 (lower ∼124 µm), while StP were found primarily in the upper third of L4 (upper ∼52 µm, Fig. 1F,G; fraction of ExN types in L4: 71% SpS, 29% StP). The distribution of interneurons (INs, Fig. 1G) was rather homogeneous over the depth of L4. At the bottom of L4, we used the steep drop of cell density from L4 to L5A as definition of the layer border (Fig. 1F,G). This included a set of apical dendrite-bearing neurons at the bottom of the thus-defined L4 which we labeled as StP, but which could also be L5A pyramidal neurons. We also found evidence that StP neurons had a higher number of somatic input synapses than SpS, yielding a slightly higher fraction of inhibitory synaptic inputs (Suppl. Fig. 3).

**Figure 1:**
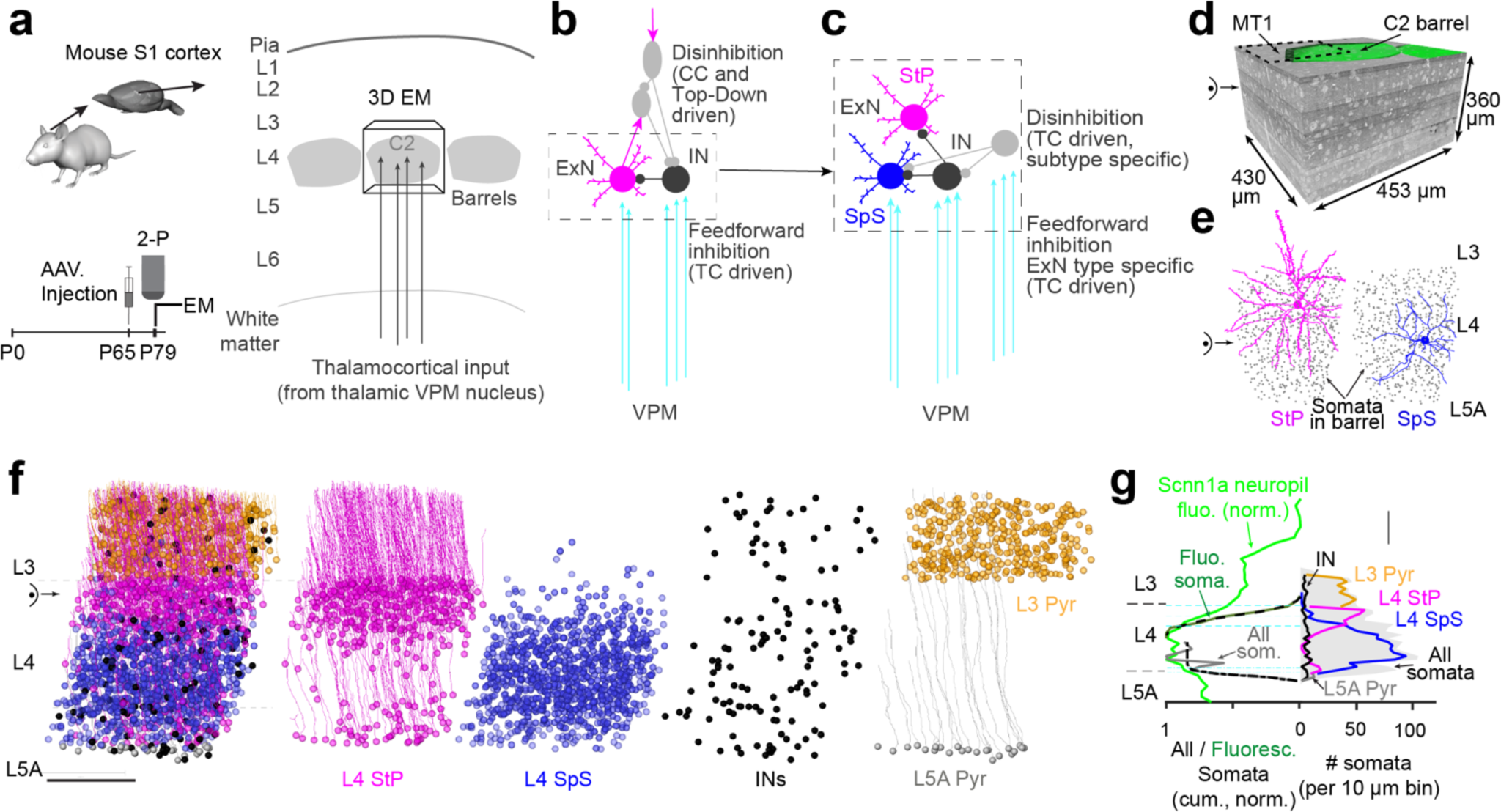
Connectomic analysis of neuronal circuitry in cortical layer 4 using 3- D electron microscopy. **(A)** Experimental design: injection of L4-targeted viral construct, functional 2-P recording experiment and 3D-EM experiment targeted to the C2 barrel in mouse S1 cortex. **(B)** Current model of thalamocortical and intracortical circuitry in which thalamic afferents drive intracortical feedforward inhibition, while intracortical disinhibitory circuits are triggered from top-down and or intracortical sources. **(C)** Circuit found in this study with thalamus-driven disinhibition on top of subtype-specific feedforward inhibition yielding a high-precision circuitry already within L4. **(D)** 3-D EM dataset spanning lower L3, L4 and upper L5A. Location of C2 barrel obtained from functional recordings superimposed (green). Note that circuit analysis was performed in a volume sized 229 x 217 x 300µm (motor tile (MT) 4) corresponding to about 1/4^th^ of the C2 barrel in the tangential plane but spanning all of L4 (dashed region). **(E)** Main excitatory cell types in L4: star pyramidal (StP) and spiny stellate (SpS) neurons. Note lack of apical dendrite for SpS cells (right). **(F)** Distribution of L4 star pyramidal (StP) neurons, spiny stellate (SpS) neurons, INs and L3 and L5A pyramidal neurons, respectively. Note slight tilt of cortical axis evident from direction of apical dendrites; data in (G) was projected onto this axis. **(G)** Distribution of cell types along cortical axis shows layering within L4 with bottom two-thirds dominated by SpS cells, top third by StP cells. Note homogenous overall distribution of INs. Definition of layer borders (dashed lines) via increase of fluorescent somata (L3->L4, green, 10^th^ percentile of sigmoid fit) and increase of soma density (L5A->L4, gray, 50^th^ percentile). Soma densities (right) shown per 10 µm along cortical axis, normalized to total number of somata. Scale bars (E, G) 100 µm.

### Thalamus-driven interneurons in L4

We next aimed to identify the thalamus-driven INs in L4 (Bagnall et al., 2011; Cruikshank et al., 2007; Porter et al., 2001; Yu et al., 2016; Yu et al., 2019). For this, we first identified all INs (Fig. 2A, n=58, of these n=52 in C2 barrel), reconstructed their dendrites (Fig. 2B, only C2 INs), and identified all input synapses made onto all their dendrites (total n=147,619 synapses onto n=52 INs, Suppl. Fig. 1; n=2,839±1,555 input synapses per IN, range 1,113-6,904 for INs with sufficiently complete dendrite, Fig. 2B-D, Suppl. Fig. 1B). We then determined for each synapse whether it was a thalamocortical synapse originating from a thalamic neuron in the ventral posteromedial nucleus (VPM) or not (Fig. 2C, total of n=2,453 VPM synapses (average of 1.6%); criteria were size and synapse number of presynaptic boutons along the putative VPM axon according to (Bopp et al., 2017; Motta et al., 2019)). This allowed a clear distinction of INs with substantially higher VPM input (n=8 of 53 INs with more than 5% VPM input synapses corresponding to at least n=98 VPM input synapses per IN, more than 3-fold the average VPM input) from all other INs which received less than 3% VPM input (Fig. 2D,E). For the remainder of the manuscript, we denote this type of INs with distinctively high VPM input as “thalamocortical interneurons” (TC INs). Interestingly, the degree of VPM input was highly distinctive also among those INs that extended a large fraction of their dendrites within the barrel (Fig. 2F,G): while all TC INs had at least 70% of their dendrites in the barrel (and received 6±1% TC input, n=8), a second group of INs with more than 70% of their dendrites in the barrel received 10- fold less TC input (0.6±0.5%, n=14, Fig. 2F).

**Figure 2:**
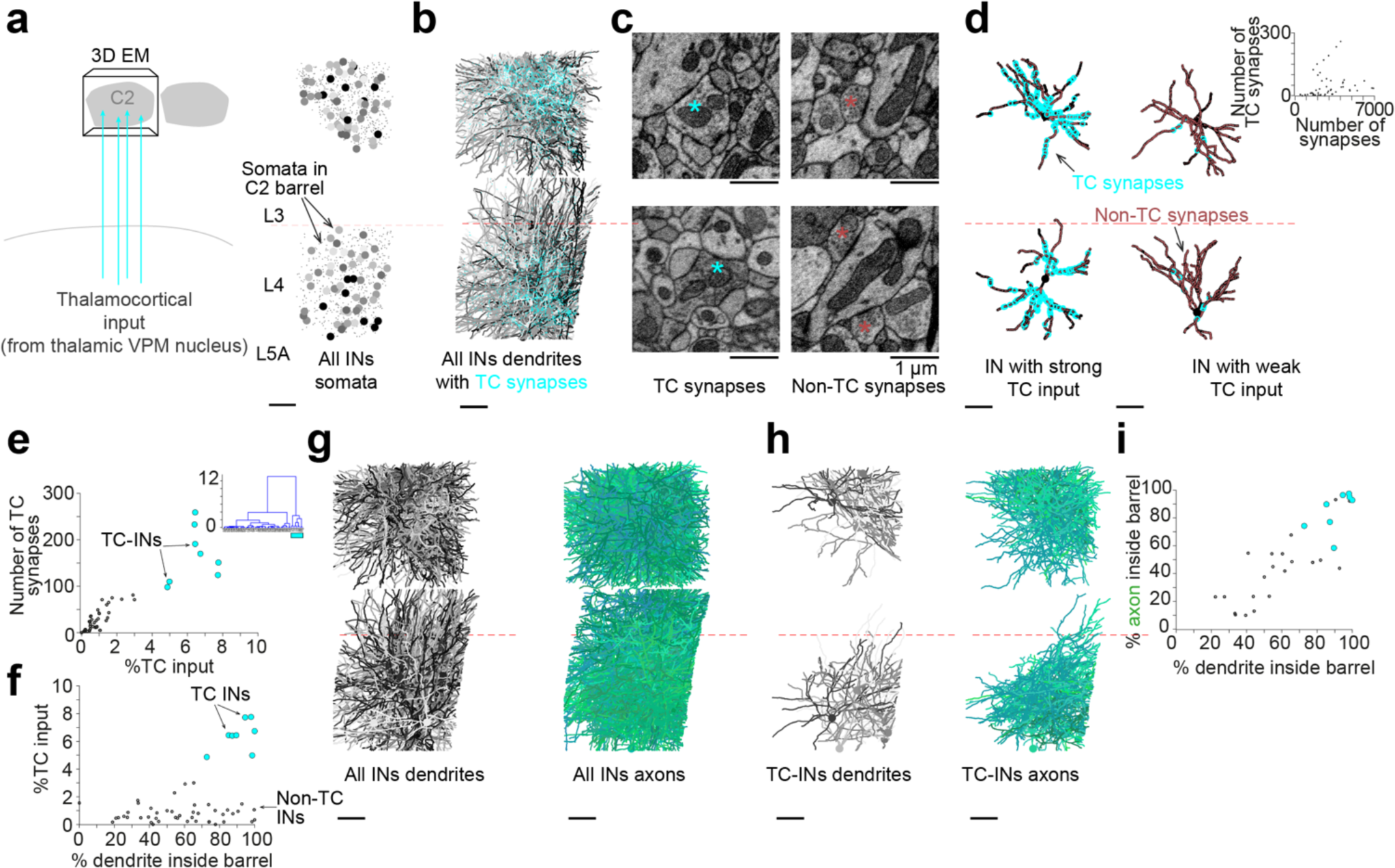
Thalamus-driven interneuron subtypes in L4. **(A)** Spatial distribution of the somata of all INs within the L4 barrel. **(B)** Reconstructions of all dendrites (grey-black) of all INs, superimposed are all n=2,453 thalamocortical (TC) synapses from the ventral posteriomedial (VPM) nucleus (cyan dots) made onto all INs. **(C)** Examples of EM cross sections showing synapses made onto IN dendrites by VPM (cyan asterisk) and non-VPM (white asterisk) boutons. TC boutons are large and make multiple spine-head synapses (Bopp et al., 2017; Motta et al., 2019) **(D)** Example dendrite reconstructions of IN subtypes receiving strong (left) and weak (right) TC input (TC synapses shown as cyan circles, non-TC synapses shown as brown dots). Inset: Distribution of all TC input synapses and all input synapses received per IN (n=53 INs). **(E)** Cluster analysis for identification of INs receiving substantial TC input (“TC INs”, shown as cyan spheres, bottom). **(F)** Fraction of TC synapses received compared to extent of dendrite path-length within the home barrel per IN. Note that even for INs with more than 70% of their dendrite extended in the L4 barrel, amount of TC input varies 10-fold between TC INs (6±1% TC input, n=8) and other INs (0.6±0.5% TC input, n=14). **(G-I)** Reconstructions of dendrites (grey-black) and axons (green) of all INs (G) compared to “TC INs” (H) indicating that TC INs also restrict their axonal output to the barrel (Koelbl et al., 2015) (I).

We then reconstructed all axons of all INs (Fig. 2G-I, Suppl. Fig. 2) and found that the TC INs all have axons that were largely restricted to the home barrel, unlike all other INs (with the exception of neurogliaform INs which had very local and barrel-related axon but received no TC input) (Fig. 2H,I). We can thus conclude that strong VPM synaptic drive converges exclusively onto barrel-related INs (Koelbl et al., 2015) whose dendrites and axons are largely restricted to the home barrel (Fig. 2I), and who constitute about 15% of all INs in L4.

### Excitatory targets of TC interneurons

Then, we analyzed the excitatory targets of the TC INs (Fig. 3). TC INs targeted somata with 23±8% of their output synapses (range 14-42%, n=8, Fig. 3A). When we identified all the ExNs innervated at their somata by TC INs (Fig. 3B), we found that some TC INs predominantly innervated SpS, while other TC INs innervated both SpS and StP, roughly according to the prevalence of these ExN subtypes in the barrel (Fig. 3C). Using this criterion, we separated the TC INs into two subtypes depending on their preference to innervate the subtypes of excitatory neurons in L4. The axonal projections of these TC IN subtypes seemed to correspond to the layered arrangement of excitatory neurons in L4: one TC IN subtype extended their axons in the lower 2/3 of the barrel, in which predominantly SpS reside (Fig. 3C, compare to Fig. 1F) while the other TC IN subtype extended their axons across the entire height of the barrel, innervating both SpS and StP. We therefore call the one subtype of TC INs lower- barrel INs (L-BINs) and the other whole-barrel INs (W-BINs).

**Figure 3:**
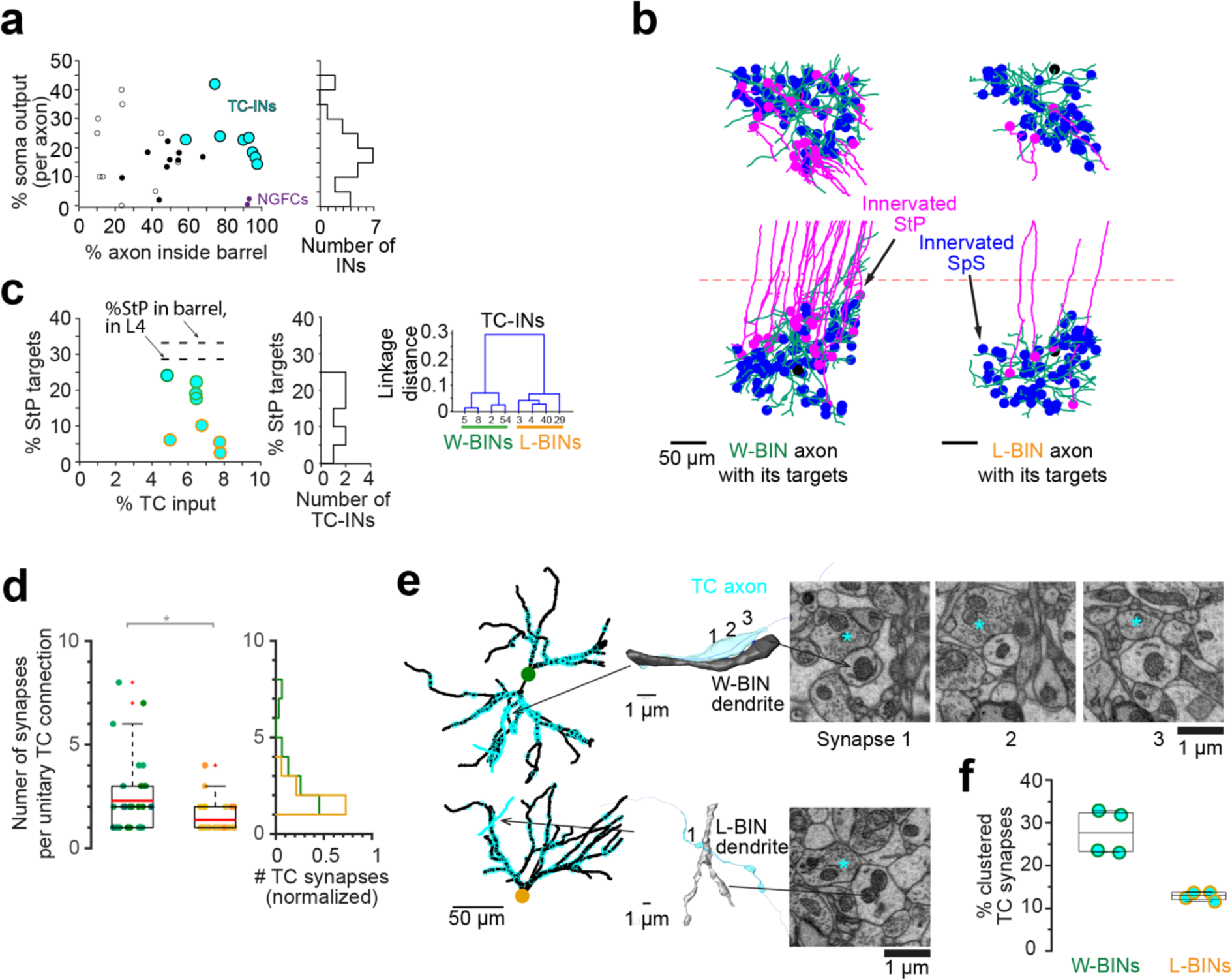
Connectomic subtypes of TC interneurons in L4. **(A)** Target preference of L4 INs to innervate somata of excitatory neurons. Note that all TC INs make 15-40% of their output synapses onto cell bodies (1420 of 6839 output synapses). 2 barrel- restricted INs with no TC input and no soma targeting were identified as neurogliaform cells (NGFCs; dark purple spheres, see Suppl. Fig. 1). Full circles, data from fully annotated IN axons; open circles, data from randomly sampled axonal output synapses of INs. **(B)** Examples of 2 TC INs with all innervated postsynaptic ExN somata shown (blue: spiny stellate somata; magenta: star pyramidal somata shown with their apical dendrite). Note innervation of both SpS and StP for TC IN shown on the left, while strong bias for innervation of SpS for TC IN shown on the right. **(C)** Cluster analysis used to distinguish subtypes of TC INs based on their innervation of postsynaptic ExN types: first group that innervate both StP and SpS somata (“W-BINs”, green circles, top), close to a homogeneous sampling of available ExN types, and second group that suppress StP innervation 3.5-fold (21±3%, N=4 vs 6±3%, N=4, p=0.03, Wilcoxon rank-sum test) and predominantly innervate SpS somata (“L-BINs”, orange circles, bottom). Note this preferential innervation goes along with a height-restricted axon for L-BINs (Suppl. Fig. 2C). **(D-F)** Analysis of TC input onto W-BINs and L-BINs shows larger number of synapses per TC axon-to-W-BIN connection (D, 2.3±1.8 TC synapses per TC axon-to W-BIN pair vs 1.4±0.7 TC synapses per TC axon- to L-BIN pair (n=31 and n=33 connections, respectively, p<0.02, Wilcoxon rank-sum test) and (E) more synapse clustering (top, 3 consecutive synapses by same TC axon) compared to single synaptic innervations (bottom). Quantification (F) shows about 2- fold difference in degree of synaptic clustering of TC input onto W-BINs vs. L-BINs (28±5% of TC innervations, N=4 vs 13±1%, N=4, p=0.029 Wilcoxon rank-sum test). Note that this difference in strength and clustering of TC input was not used for the definition of L-BINs vs W-BINs in a-c.

### Thalamic input to TC interneurons

When quantifying the TC input onto W-BINs and L-BINs, we found that W-BINs received 2.3±1.8 TC synapses per TC axon-to IN pair (evaluated using 20 fully reconstructed TC axons), while L-BINs received 1.4±0.7 TC synapses per axon per cell (n=31 and n=33 connections, respectively, p<0.02, Wilcoxon rank-sum test, Fig. 3D). While mapping these TC input synapses to the L-BINs and W-BINs, we had further noticed that in some cases, TC axons made several synapses onto the same IN dendrite in close proximity (Fig. 3E, see (Bagnall et al., 2011; Cruikshank et al., 2007; Porter et al., 2001; Sun et al., 2006; Yu et al., 2016)). When quantifying the fraction of TC input synapses made in such a clustered configuration, we found a clear difference between the W-BINs (with 28±5% clustered TC synapses) and L-BINs with 2-fold less clustering (13±1%, p=0.029, Wilcoxon rank-sum test, Fig. 3F).

Since this difference in strength and clustering of TC synaptic innervation had not been used for defining the subtypes of W-BINs vs. L-BINs, it already served as a first indication of systematic connectomic difference of these IN subtypes.

### Interneuron-to-Interneuron connectivity

We then wanted to understand the connectivity between interneurons within the barrel. For this, we identified all contacts between all IN axons (n=30 INs with at least 1.5mm axonal path length reconstructed, Suppl. Fig. 2) and all IN somata and dendrites and determined whether there was a chemical synapse between them (Fig. 4A,B). The resulting 30x43 IN-to-IN connectome (Fig. 4B) showed that (1) already among the TC- INs, substantial disinhibitory circuitry exists (Fig. 4C); (2) the thalamus-driven disinhibitory circuit is strongly directed from L-BINs to W-BINs with 10-fold less innervation in the opposite direction (3.8±4.1 synapses per connection vs 0.38±0.89 synapses per connection, n=16, p=0.002, Wilcoxon rank-sum test, Fig. 4D,E); (3) L- BINs receive least inhibition from any other of the INs (Fig. 4F).

**Figure 4:**
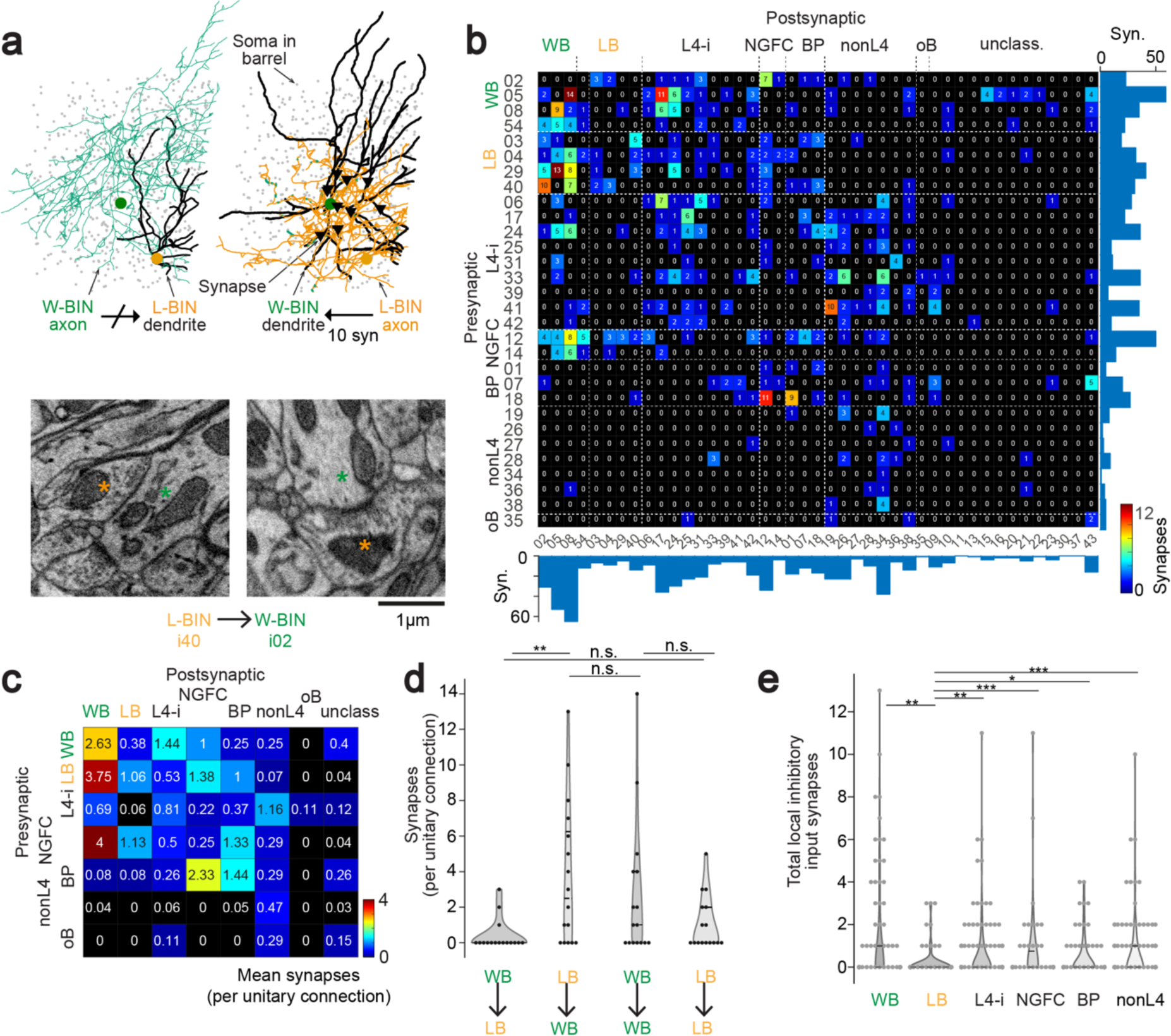
IN-to-IN connectome in L4. **(A)** Example reconstructions of pair of W-BIN (i02) and L-BIN (i40) showing inhibitory synapses (black triangles): no inhibitory synapse in the W-BIN-to-L-BIN connection (left), but n=10 synapses in the reciprocal connection (right). Bottom: Examples of EM cross sections showing synapses made by L-BIN axon (orange asterisk) onto W-BIN dendrite (green asterisk). **(B)** 30x43 IN- to-IN connectome; numbers in connectome represent synapses per directed connection. Zeros mean no synapse found after examination of all outgoing connections. Only INs with at least 1.5 mm axon length shown as presynaptic (see IN gallery in Suppl. Fig. 2; row/column indices correspond to IN IDs in gallery). L4-I, L4- related IN; NGFC, Neurogliaform cells; BP, Bipolar INs, nonL4, non-layer related INs; oB, neighboring barrel IN, unclass, INs with less than 1.5 mm axon (see Suppl. Table 1) **(C)** 7x8 IN type- to IN type connectome, from aggregation of blocks in (B). Note directed L-BIN-to-W-BIN inhibition that is about 10-fold stronger than inverse direction, see (d). Numbers in connectome indicate average number of synapses per pair of pre-and postsynaptic INs. **(D)** Comparison of number of synapses per L-BIN-to-W-BIN connection vs. the reverse direction (Wilcoxon rank-sum test, p<0.0001) and within each group (W-BIN to W-BIN vs L-BIN to L-BIN, p=0.32, Wilcoxon rank-sum test). **(E)** Total inhibitory input synapses per INs of each subtype, aggregated over all inputs from INs of other subtypes. Inhibition onto L-BINs was significantly lower than inhibition received by other IN subtypes (W-BINs vs L-BINs p<0.001, L4-INs vs L-BINs p<0.01, NGFCs vs L-BINs p<0.001, Bipolar INs vs L-BINs p<0.05, nonL4 INs vs L-BINs p<0.001, Wilcoxon rank-sum test).

### Possibility of gap-junctional coupling

In addition to chemical synaptic connectivity, L4 INs were shown to exhibit electrical synaptic coupling during development (Beierlein et al., 2000, 2003; Connors, 2017; Galarreta and Hestrin, 1999; Gibson et al., 1999; Hatch et al., 2017; Hestrin and Galarreta, 2005; Pernelle et al., 2018). While the direct detection of gap junctions is not possible in EM data of the utilized resolution, a necessary condition for gap junctional coupling between neurons is that their dendrites (or somata) form direct appositions (without glia or other neurons in between, Fig. 5A). We therefore analyzed the degree of direct dendro-dendritic touch between all interneurons as an upper bound on the number of appositions per IN pair at which gap junctions could in principle be formed (Fig. 5B). While IN dendrites did form direct appositions, only 3% (n=31 of 946) of IN pairs had more than one apposition, and 87% (n=822 of 946) had no apposition at all, excluding gap junctional coupling in these IN pairs. When analyzing the dendro-dendritic touches for IN subtypes (Fig. 5C), we found that L-BINs (n=4/6 pairs) were the only IN types in which a majority of IN pairs within the type formed direct dendro-dendritic touches. No other IN type or combination of IN types did so (only types with at least 4 INs analyzed). Since the degree of gap junctional coupling among INs in L4 had been reported to occur between about 2/3-3/4 of the INs (Beierlein et al., 2000, 2003; Gibson et al., 1999), L-BINs are a possible candidate for such coupling. For other IN types, 56-100% of INs within these groups had no dendro- dendritic touch (Fig. 5D), excluding the possibility that strong and prevalent electrical coupling exists among these other IN neuron populations. Notably, when analyzing all TC INs together (combination of W-BIN and L-BIN), only a minority of IN pairs formed direct appositions (n=13 of 28), further indicating the relevance of the differentiation between L-BINs and W-BINs.

**Figure 5.**
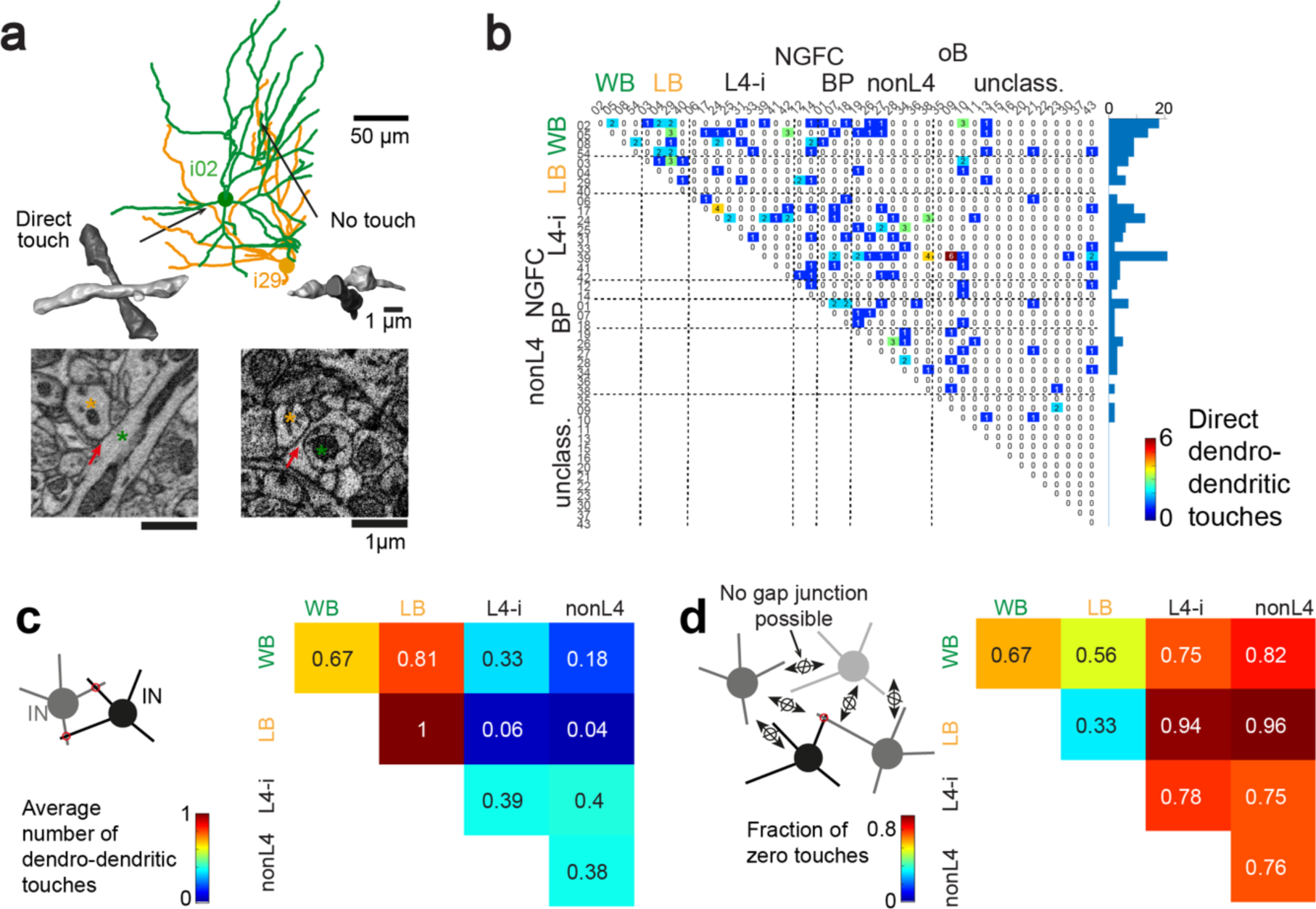
Dendrodendritic touch as prerequisite of gap junctional coupling. **(A)** Example dendrite reconstructions of IN pair for which 2 apparent dendro-dendritic touches were found (dendrites closer than 1 µm). EM cross sections (bottom) showing that one of these corresponded to a direct dendritic touch (left), while the other dendritic location was separated by a thin glial sheath (bottom right), thus prohibiting the existence of gap junctional coupling at this location. **(B)** Frequency of direct dendro- dendritic touch between IN dendrites (dendrite contactome). Note dominance of zero or one-contact cell pairs (97%, n=915 of 946). **(C)** Same as (B) aggregated for IN types reporting the average number of dendro-dendritic touches per IN cell pair within and across IN types. Only IN types with at least 4 INs shown. **(D)** Fraction of IN cell pairs for which no dendro-dendritic contact was found. Note that only for L-BINs, a majority of IN cell pairs within each type had at least one touch. In all other IN types, gap junctional coupling cannot occur in a majority of within-type IN pairs (Beierlein et al., 2000; Gibson et al., 1999).

### Summary of connectomic IN types

To recapitulate, the finding of a directed chemical disinhibitory circuit between L-BINs and W-BINs was surprising for two reasons: first, disinhibitory circuitry has so far been proposed to be involved in intracortical (Gainey et al., 2018; Karnani et al., 2016; Pfeffer et al., 2013; Rikhye et al., 2021; Xu et al., 2013; Yu et al., 2016; Yu et al., 2019) and top-down (Lee et al., 2013; Williams and Holtmaat, 2019) processing, but not for the earliest stage of cortical activation from the thalamus. Secondly, the subtype specificity of this circuitry implies a relation between the excitatory output selectivity and disinhibitory connectivity: those TC-INs that innervate all types of excitatory neurons in the barrel (W-BINs) are unidirectionally inhibited by those TC-INs that preferentially target spiny stellates. Furthermore, the lack of disinhibitory input onto L- BINs but not W-BINs sets the L-BINs apart as a strong inhibitory circuit component, that itself does not receive substantial inhibition. Rather, the data on dendro-dendritic touch may indicate that L-BINs correspond to INs that were reported to be electrically coupled among each other (Beierlein et al., 2000, 2003; Gibson et al., 1999).

### Sequence of thalamocortical innervation

We next investigated the TC input axons converging on the L4 circuit in more detail from the TC axon perspective (Fig. 6). For this, we reconstructed 20 TC axons (identified as originating in the VPM nucleus by their trajectory from the white matter, myelination until entering L4, and features of axonal boutons (see (Bopp et al., 2017) and (Motta et al., 2019), Suppl. Fig. 4A)) and identified all of their output synapses and the types of synaptic targets (Fig. 6A-C). TC axons (n=20 axons, total length: 16.22 mm, synapse density: 0.23±0.05 per µm; 3,566 output synapses total) made 4±2 % (n=20) of their synapses onto the shafts of smooth dendrites, all others (96%) were onto spines of spiny dendrites. TC axons and their synapses yielded a substantial gradient of TC innervation from lower to upper L4 (Fig. 6c, Suppl. Fig. 4B see ref. (Motta et al., 2019; Oberlaender et al., 2012)).

**Figure 6:**
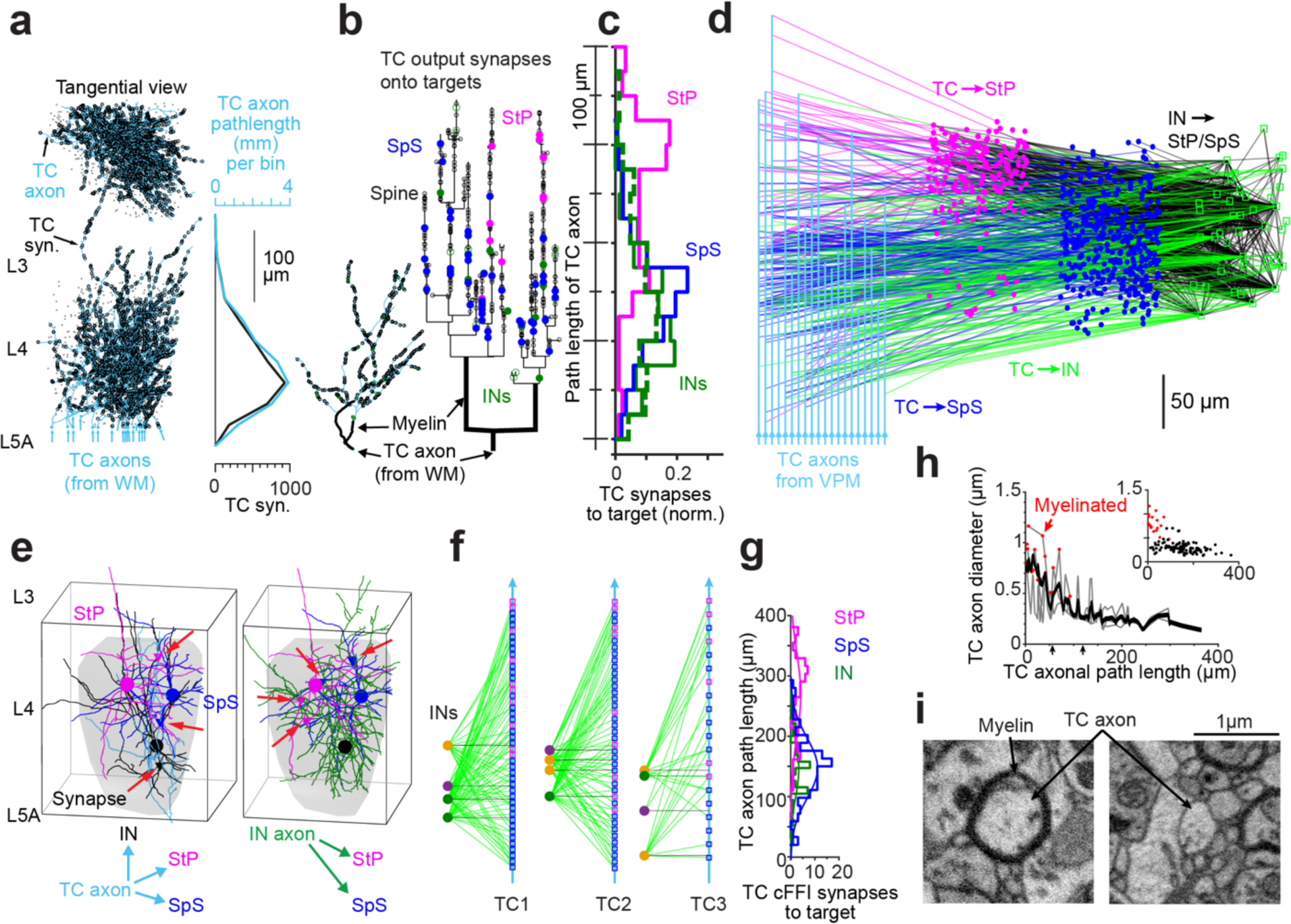
Thalamocortical innervation, cellular feedforward inhibition and axonal synapse sorting. **(A)** Projection of 20 TC axons into the barrel and distribution of TC path length and synapses along cortical axis, yielding a substantial TC synaptic gradient in upper L4, see (Motta et al., 2019) and Suppl. Fig. 4 for TC definition. **(B)** Position of output synapses along one TC axon shown in axonogram as function of axonal path length (inset: single TC axon reconstruction). TC synapses onto INs (green; open circles, smooth dendrites; filled circles, identified INs) and subtypes of excitatory neurons (StP, magenta; SpS, blue) showing synaptic sorting according to postsynaptic cell type. **(C)** Ensemble distribution of output synapses from n=20 TC axons. Note overlap of TC-to-IN with TC-to-SpS innervation, offset from TC-to-StP innervation (offset of peaks, 117 µm between IN and StP vs 17 µm between IN and SpS). Dashed green: output to smooth (IN) dendrites without identified IN soma in barrel. **(D)** Circuit diagram between 20 thalamocortical axons, 43 INs, 375 StP and 964 SpS targets (3089 synapses total) indicating feedforward-inhibition (FFI) circuits with differential TC synapse positions. **(E)** Cellular-level precision of FFI-triads (red arrows indicate synapse positions) in TC–to-L4 circuits. Gray shading corresponds to barrel volume as in Fig. 2. **(F)** INs (circles, left) innervated by a given TC axon (cyan, black lines indicate first TC-to-IN synapse along the TC axon) innervate 96% of the SpS (blue squares) and 93% of StP (magenta squares, IN synapses as green lines) that were also innervated by that TC axon (three fully reconstructed TC axon examples; 100% ExN targets match between IN and TC synapses when only considering StP and SpS within the barrel hollow). **(G)** Histogram of TC synapse positions for the cFFI triads, indicating coincident activation of IN and SpS, offset from activation of StP. **(H)** TC axon diameters show substantial reduction in the unmyelinated TC axon within L4, making timing effects of synapse sorting plausible (compare (Schmidt et al., 2017)). Axon diameter was measured at random locations along 3 TC axons (grey lines, average, black, single measurements in inset). **(I)** Example cross-sections of TC axon at 44 µm (left) and 122 µm (right) path length showing difference in myelinisation and diameter. Scale bars, 100 µm (A-C); 50 µm (D); 1µm (I).

When differentiating the TC innervation of SpS and StP (Fig. 6C), SpS and StP did not differ in the number of TC input synapses they received (1.18±0.50 synapses (n=22) per TC-to-StP connection versus 1.33±0.83 (n=27) per TC-to-SpS connection, Wilcoxon rank-sum test, p=0.47, Fig. 6A,B). Synapse size did also not differ between TC synapses onto SpS and StP (p>0.05, n=17 StP, n=40 SpS, Suppl. Fig. 4C,D).

We found, however, that TC axons sequentially activate first SpS and then StP. The peak of these innervations is offset by at least 100 µm along the TC axon (Fig. 6C, pooled innervations; for detailed analysis, see below). Notably, the TC innervation of interneurons aligned with the position of TC synapses onto SpS (Fig. 6C, offset less than 20 µm). With this, we found that the layering of excitatory cell bodies in L4 yielded a sequential TC activation of these excitatory neuron populations, in addition to the differential innervation by subtypes of interneurons (Fig. 2).

We then investigated how the inhibitory neurons innervated by TC axons projected back into the excitatory circuit in L4 (Fig. 6D-G). Feedforward inhibition (FFI) has been described as a hallmark of TC processing (Agmon and Connors, 1991; Bruno and Sakmann, 2006; Bruno and Simons, 2002; Gabernet et al., 2005; Porter et al., 2001; Temereanca and Simons, 2004; Torii and Levitt, 2005; Yu et al., 2016; Yu et al., 2019) but experiments were so far not able to discern whether this FFI was a population effect or whether it operated at the precision of cellular FFI (cFFI (Schmidt et al., 2017)). Here, we found the cFFI configuration in at least 95% of TC-to-ExN connections were matched by at least one cFFI connection (evaluated in detail for three TC axons, Fig. 6F; n=135 TC-to-ExN synapses, n=17 TC-to-IN synapses, n=368 IN-to-ExN synapses with 103 of 108 excitatory targets; see Methods). Notably, for these cFFI configurations, the TC synapses innervating the INs were spatially aligned with the output to SpS, but 122 µm offset from the synapses onto StP, which were also innervated by cFFI (Fig. 6F,G; Wilcoxon rank-sum test p<10^-10^ for output to INs vs. StP; p<10^-18^ for SpS vs. StP; position of output to INs and SpS was indistinguishable, Wilcoxon rank-sum test, p>0.9). The previous results from cFFI in rat medial entorhinal cortex (Schmidt et al., 2017) had indicated that such a sorted axonal innervation could efficiently control postsynaptic excitatory activity if the presynaptic excitatory axon was small and had moderate action potential (AP) conduction velocity. For the TC axons, after myelination ended, we found diameters of 0.29±0.10 µm (mean±s.d., n=3) within L4 (Fig. 6H,I). This would indicate a higher conduction velocity, such that conduction-based delays may be smaller than in MEC (Schmidt et al., 2017).

### Circuit-based prediction of differential L4 activation

Our key circuit findings in L4, the direct TC-driven disinhibitory circuitry with selective inhibition of SpS vs. StP targets, would allow the following predictions about functional activation in L4 by sensory stimuli (Fig. 7). In the case that only W-BINs but not L-BINs become activated by TC input (which could be plausible via the highly clustered TC input to W-BINs, but not L-BINs, Fig. 7A), a narrow window of activity would be predicted both in SpS and StP targets, and the activity of StP could even be lower or shorter than SpS if the conduction delay effects matter in the circuit (Fig. 7A). In case of selective TC activation of L-BINs (Fig. 7B), the lack of cFFI onto StP would predict a broad window of activation, potentially with very strong responses, in StP, while SpS would remain briefly activatable, only, given the selective cFFI of L-BINs onto SpS, but not StP.

**Figure 7.**
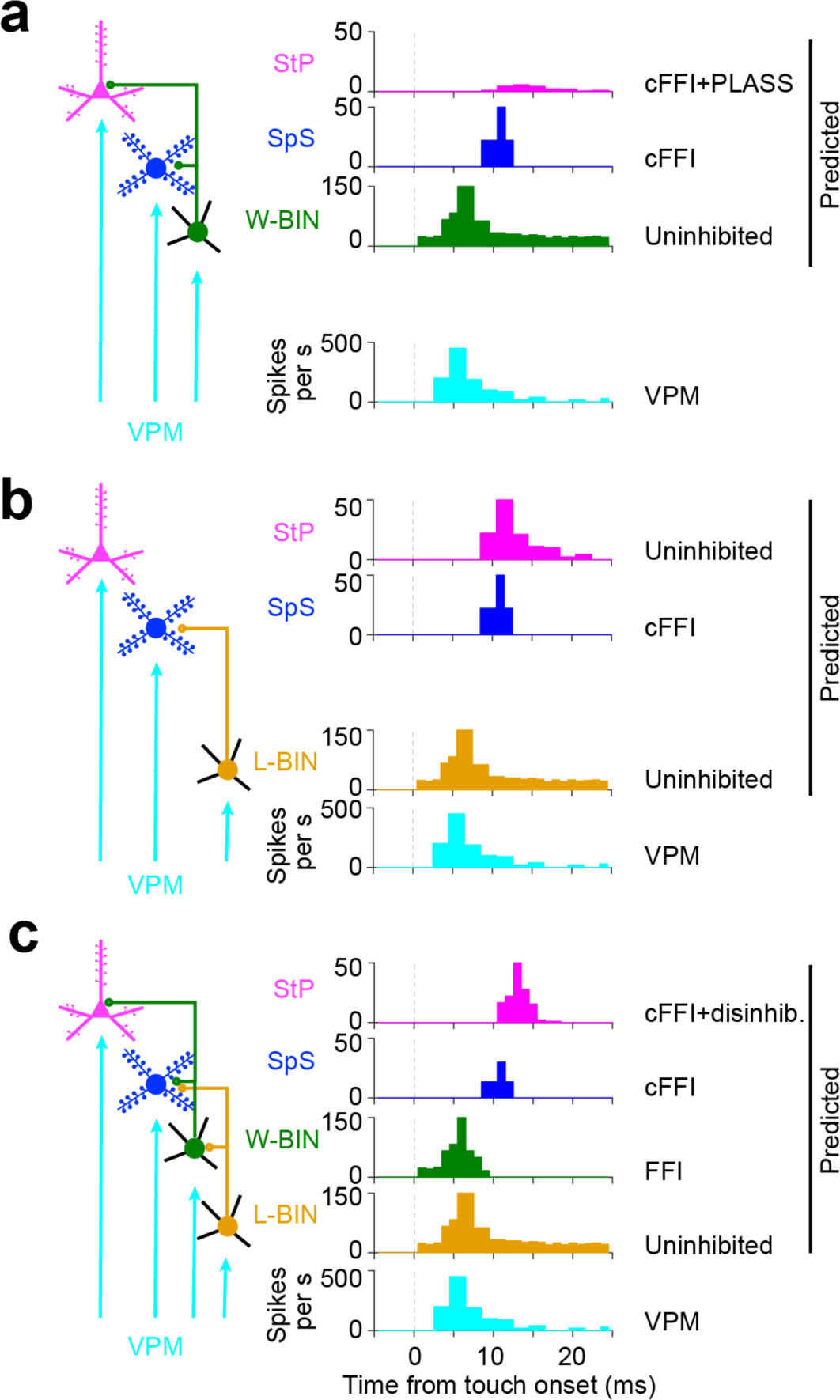
Possible functional effects of TC-driven subtype-specific disinhibitory circuitry in L4. **(A)** TC activation of ExNs and exclusive activation of W-BINs would yield cellular feedforward inhibition, effectively increasing activation threshold in ExNs(Yu et al., 2016; Yu et al., 2019) and limiting the time window of activation for SpS and StP. The synaptic offsets in the W-BIN -to-StP pathway could even suppress StP activity (as in (Schmidt et al., 2017)). **(B)** Exclusive TC activation of L-BINs but not W-BINs would yield cFFI for SpS (as in a), but would leave StP without direct thalamus-driven FFI, thus with potentially strong and broad StP activity. **(C)** Activation of the entire TC driven circuitry in L4 would initially yield cFFI onto both SpS and StP, which would be stronger on SpS (two input IN populations). Subsequently, the L-BIN-to-W- BIN inhibitory connectivity could curtail W-BIN activity, thereby relieving StP from cFFI, yielding a stronger StP response. (A) would be plausible based on the strong TC clustering for W-BINs, which could allow a selective activation of W-BIN but not L-BIN for sparse or weak thalamic input. Strong thalamic activity could yield setting (C). Note further that instead of impulse response, these circuits could yield differential frequency responses to various temporal patterns of whisker inputs ((Arabzadeh et al., 2005; Isett et al., 2018; Jadhav et al., 2009; Wolfe et al., 2008) see Discussion).

In case of combined activation of L-BINs and W-BINs, however (Fig. 7C), the specific circuitry in L4 would predict a brief activation in SpS that is curtailed strong cFFI from both L-BINs and W-BINs. Similarly, StP would initially receive strong cFFI, potentially supressing immediate activation. After strong activity of L-BINs has occurred, however, the targeted inhibitory circuit from L-BINs to W-BINs would predict a strong reduction in W-BIN activity. With this, the cFFI onto StP would be lifted, and StP could show strong activation by its TC input. This would yield a selective activation of L4 ExNs: spiny stellates would only briefly activate, but StP would become strongly active after a brief delay of potentially a few milliseconds.

## DISCUSSION

Our analysis of the thalamus-driven inhibitory circuitry in cortical layer 4 revealed (1) Highly specific VPM innervation of about 15% of interneurons in the barrel yielding a clear distinction of TC-driven INs (TC-INs); (2) All TC-INs have a barrel-restricted axonal projection (synaptic output); (3) TC-INs fall into two classes based on their innervation of spiny stellate vs. star pyramidal cells, implemented via the strong layering of these target cells within L4; (4) Among the TC-INs, a directed disinhibitory circuit exists from lower-barrel INs (L-BINs) to whole-barrel INs (W-BINs); (5) the thus defined IN types differ in the strength and degree of clustering of the TC synaptic input they receive; (6) thalamus-driven FFI circuitry is configured at cellular precision (cFFI), with sorted synaptic outputs along TC axons.

Together, these findings describe a highly precise thalamus driven inhibitory circuitry that would be capable of creating differential temporal windows for activation of the subtypes of excitatory neurons within L4, possibly allowing for differential representation of sensory stimuli of varying temporal structure.

### Connectomic definition of IN types

We used exclusively connectomic information for the distinction of types of INs in L4: strong TC innervation and synaptic target preference for excitatory subtypes for the initial IN type definition; and then found further connectomic specializations of these IN subtypes in their directed disinhibitory synaptic connectivity, and strength and clustering of TC input. It was notable that these connectomic differences were multi- fold: 6-fold stronger TC innervation (Fig. 2F); 3.5-fold difference in ExN targeting (Fig. 3C); 10-fold bias in disinhibitory connectivity (Fig. 4C); 1.7-fold difference in unitary TC input synapse number (Fig. 3d) and 2.2-fold difference in TC clustering (Fig. 3F). Such clear connectomic patterns may be promising for approaching the still debated field of IN type definition (Ascoli et al., 2008; Yuste et al., 2020) from a connectomic perspective (as exemplified before in the retina (Helmstaedter et al., 2013)).

### Relation to SST IN types

Recent work (Yu et al., 2019) has described the differential TC activation of subtypes of INs identified via their protein expression (parvalbumin (PV) *vs*. somatostatin (SST) and vasointestinal peptide (VIP) (Pfeffer et al., 2013; Rudy et al., 2011; Scala et al., 2019; Taniguchi et al., 2011; Tremblay et al., 2016)). In this data, disinhibition is implied to be provided by SST and VIP INs, while PV-positive INs are the only INs to receive short-latency TC activation (Yu et al., 2019). Our data indicates that even subtypes of locally projecting soma-targeting INs (likely identified with PV-positive INs) differentially provide disinhibition within the barrel (Fig. 4). Since SST-INs are described as preferentially dendrite-targeting (Pfeffer et al., 2013; Scala et al., 2019; Williams and Holtmaat, 2019), it is unlikely that the L-BINs correspond to these IN subtypes. Rather, the identification by the classical IN markers may not be sufficient in L4. Recent data has proposed that SST INs have unconventional morphology in S1 (Scala et al., 2019; Zhou et al., 2020). It has to be determined whether the circuits we found are related to the conventional expression markers widely employed (see also ref. (Koelbl et al., 2015) for the (lack of) relation between morphology, markers and electrophysiology in L4, and Suppl. Table 1 for a tentative comparison of our INs to published data).

### Sublayers within cortical layer 4 in other species

In primates, cats and higher mammals (Braak, 1976; Clark, 1925; Collins et al., 2005; Hassler and Wagner, 1965; Kaas et al., 1972; Lund, 1973; Otsuka and Hassler, 1962; Wong and Kaas, 2008, 2009, 2010) cortical L4 in primary sensory areas has been subdivided since the initial histological analyses (Brodmann, 1909; Ramón y Cajal, 1899). These subdivisions were based on overall histoarchitectonic differences in cell body density (Brodmann, 1909; Campbell et al., 1905; Garey and Harris, 1971; Lewis and Ferrier, 1880; Ramón y Cajal, 1899; Valverde, 1977; von Bonin, 1942). In macaque and cat, the observation of more pyramidal cells towards L3 gave rise to a dispute of where to define the upper border of L4 (Garey and Harris, 1971). A subdivision of L4 by excitatory cell type was not clearly accepted in these species. For rodents, a subdivision of L4 has not been described, and even less one related to the types of excitatory cells – yet, with the benefit of hindsight, some evidence can be found (Simons and Woolsey, 1984a). Since the stacked positioning of neurons that we find (Fig. 1) is only clearly visible within the barrel, but not in the septa, and since it occurs on spatial scales of less than 200 µm, it may have evaded light-microscopy-based circuit analysis.

### cFFI in mammalian cortex

While feedforward inhibition as a key wiring principle has been identified in many mammalian circuits in the cortex (Alger and Nicoll, 1982; Bruno and Simons, 2002; Buzsáki, 1984; Cruikshank et al., 2007; Duguid et al., 2015; Isaacson and Scanziani, 2011; Kanichay and Silver, 2008; Mittmann et al., 2005; Pouille and Scanziani, 2001), it was not possible to distinguish between population-level or cellular-precision FFI. 3D EM has allowed the systematic investigation of this, and so far has identified cFFI as a prevailing wiring principle intracortically in supragranular layers (Schmidt et al., 2017) and for TC input to cortex (this study, Fig. 3). In combination with axonal synapse sorting (PLASS (Schmidt et al., 2017)), such circuits can theoretically very precisely control the propagation of excitatory activity (Schmidt et al., 2017). While synaptic placement as a means for temporally precise neuronal activity has been long known in non-mammalian species (Carr and Konishi, 1990; Kornfeld et al., 2017) the degree of precision in mammalian cortical tissue is surprising. Whether AP conduction in TC axons is slow enough to allow full excitatory suppression (Schmidt et al., 2017) or rather a narrowed window of postsynaptic activation will have to be investigated. The level of circuit precision observed here indicates that very detailed studies are required for an analysis of the capacity of these circuits in mammals.

### Circuit effects on temporal patterns of sensory stimuli

The surprising circuit-level differences in the thalamic feedforward convergence onto spiny stellates vs. star pyramidal neurons (Fig. 7) could allow for differential representation of sensory stimuli with different temporal structure. In particular upon strong VPM activation, when both W-BINs and L-BINs are predicted to be active (Fig. 7C), spiny stellates are predicted to respond with higher temporal fidelity to VPM input, while StPs are predicted to have a delayed response governed by the disinhibitory delay (the time it takes to inhibit W-BINs by L-BINs), that then however could be much stronger than in SpS. In principle, such a configuration could imply a high-vs low-pass filtering of sensory inputs. One can speculate that this could correspond to the representation of smooth (high temporal input frequency) vs. rougher (low temporal frequencies) surfaces via the principal whisker (Arabzadeh et al., 2005; Carvell and Simons, 1990; Isett et al., 2018; Jadhav et al., 2009; Park et al., 2019; Wu et al., 2013). Key caveats to this interpretation are whether the L-BIN connection to W-BINs is strong enough to fully suppress W-BIN activity even under concomitant thalamic input; and whether other inhibitory sources may become important already within the first 10 ms after the sensory stimulus.

### Outlook

We have provided a first connectomic description of the thalamus driven inhibitory L4 circuits in mammalian sensory cortex. Future work, that should include the structure of the excitatory intracortical networks, will reveal whether the level of precision found here in synaptic and cellular architecture is an overarching principle of mammalian cortex.

## METHODS

### Animal experiments

All experimental procedures were performed according to the law of animal experimentation issued by the German Federal Government under the supervision of local ethics committees and according to the guidelines of the Max Planck Society. Scnn1a-Tg3-Cre mice were purchased from Jackson Laboratory (US) and bred in animal facility at CAESAR institute (Protocol 84-02.04.2014.A331 from the Landesamt für Natur, Umwelt und Verbraucherschutz, Nordrhein-Westfalen, Germany).

### Viral injections

The AAV virus (AAV9.CAG.Flex.GCaMP6s.WPRE.SV40) injection was performed on male animals ranging in age from 56 to 72 days. Anesthesia was induced with ketamine and xylazine (120 and 5 mg per kg of body weight, respectively) and maintained using supplementary doses between 10 and 20 % of the induction dose whenever the animal began to recover withdrawal reflexes. Through all procedures, body temperature was maintained at around 37 °C using a regulated heating blanket and rectal thermal probe. A small craniotomy (approximately 0.5 mm × 0.5 mm) was created at the location of 1.45 mm caudal from bregma and right lateral 3.80 mm, which was 1 mm lateral to the center of the whisker representation in the somatosensory cortex. After removal of dura mater, a log-taper glass pipette (tip beveled at 30°, inner diameter 8-15 μm) filled with the virus stock solution was advanced 1.1 mm towards the midline from cortical surface at an angle of 15° from the horizontal, placing the pipette tip at a depth of 650 μm in the target area. 200 μl virus solution was injected over 5 min. After retraction of the pipette, the skin incision was then closed with Vicryl sutures and the animals were allowed to recover. Expression time ranged from 14 to 21 days. On the day of Ca^2+^ imaging experiment, animals were again anesthetized with ketamine and xylazine, the skull over the target area (1.45 mm caudal from bregma and right lateral 2.80 mm) was exposed, and a head plate for the stabilization of the head under the microscope was attached. A large craniotomy (approximately 2.5 mm × 2.5 mm) was made, centered on the target area, and the dura mater was opened. Astrocytes were counterstained by applying sulforhodamine 101 briefly to the brain surface and the preparation was stabilized using 1% agarose (Sigma-Aldrich, US, dissolved in artificial cerebrospinal fluid of following composition in mM: NaCl, 135; KCl, 5.4; CaCl2, 1.8; MgCl2, 1; HEPES, 5) and a coverslip to suppress brain movement during multiphoton calcium imaging.

### In vivo two-photon calcium imaging with whisker stimulation

Labeled neurons and astrocytes were visualized using a custom-built multiphoton microscope(Ramachandra et al., 2020). Excitation light was provided by a Ti:Sapphire pulsed laser system (Mai Tai, Spectra Physics, CA, USA) tuned to 920 nm. The imaging system consisted of a galvanometric system to generate fast scanning and a 20x objective lens (WPlan-APOCHROMAT, Carl Zeiss, Germany). The barrel field neuropil imaging was carried out at 256 x 256 pixel resolution with a frame rate of 1 Hz. The functional imaging of cell bodies was performed at 64x128 pixel resolution with a frame rate of 18.6 Hz. Motion within each frame of the imaging datasets was corrected using red fluorescence from sulforhodamine 101(Greenberg and Kerr, 2009). The single-whisker stimulation was carried out on the contralateral whisker pad using a piezo-driven plastic pipette tip to generate 15° backward and forward movement of target whisker with 0.5 second interval. The stimulation was repeated every 8 seconds. At the end of the imaging section, a 2-channel image stack was acquired to register positions of fluorescent cells. The stack contains 600 slices starting from pia towards white matter at a step size of 1 μm and covering an area of 300 × 300 μm^2^ at a pixel size of (1,172 nm)^2^ in-plane.

### Histology

At the end of the imaging experiment, the animals were perfused transcardially with 30 ml 0.15 M cacodylate buffer, following at least 90 ml of fixation solution (2.5 % paraformaldehyde, 1.25 % glutaraldehyde, 2 M calcium chloride in 0.08 M cacodylate buffer, pH 7.4). After the perfusion, the animal was decapitated and the head was immersed in the same fixation solution at 4 °C overnight. For better access to the fixative, the coverslip was removed from the top of the craniotomy window. Extraction of the previously imaged cortical tissue was done on a stereotaxic instrument (Kopf) with modified ear bars to adapt to the animal head plate. A 1 mm diameter biopsy punch (KAI Medical, Honolulu, USA) mounted on an electrode manipulator was centered on top of the target tissue and advanced along cortical axis from cortical surface towards white matter for about 2 mm to cut out the target tissue for *en bloc* EM staining.

### Sample preparation for SBEM

*En bloc* staining for SBEM was performed as in ref.(Hua et al., 2015) with minor modification. Briefly, the tissue was stained with an osmium tetroxide solution (2 % OsO4 in 0.15 M cacodylate buffer, pH 7.4) followed by incubation with 2.5 % ferrocyanide (in 0.15 M cacodylate buffer, pH 7.4) for 90 min and an additional incubation with 2 % OsO4 solution (in 0.15 M cacodylate buffer, pH 7.4) for 45 min. Subsequently, the sample was incubated in saturated aqueous thiocarbohydrazide (TCH) solution for 60 min, 2 % OsO4 aqueous solution for 90 min at room temperature and incubated in unbuffered 1 % uranyl acetate (UA) aqueous solution at 4°C overnight. On the next day, the sample was incubated for 2 hours at 50 °C in 1 % UA aqueous solution and 0.03 M lead aspartate solution (pH 5.0). Dehydration and resin embedding were performed with ethanol and acetone, infiltrated with 1:1 acetone and Spurr’s resin (Sigma) mixture at room temperature overnight and then with pure resin for 6 hours before baking in a pre-warmed oven at 70 °C for 3 days.

### Cortical vascular imaging and global alignment

The embedded sample was mounted on an aluminum pin along the cortical axis with conductive epoxy glue (Henkel, Germany) and then trimmed to a cube of 1×1×0.8 mm^3^. Pia matter was exposed by carefully removing a thin layer of resin with an ultramicrotome (UC7, Leica, Germany) equipped with a diamond-blade (Diatome, Switzerland). The smoothed tissue block was then coated with a 50 nm thin gold layer (ACE600, Leica, Germany), before being imaged in a field-emission SEM (Quanta 250 FEG, FEI) equipped with a custom-built in-chamber ultramicrotome (courtesy of W. Denk(Denk and Horstmann, 2004)). Using 5 keV acceleration energy and 1 μs dwell time, 538 consecutive image planes of dimensions were acquired at a pixel size of 378.9 nm × 378.9 nm and a cutting thickness of 150 nm, resulting in an image stack with a dimension of 1.164 x 0.776 x 0.096 mm^3^. To align the light-microscopic imaged C2 barrel in the sample block prepared for 3D EM, a global 3D affine alignment was performed using manually reconstructed cortical blood vessels as landmarks.

### Cortical neurite imaging, alignment and correspondence

The sample was then imaged using a field-emission SEM (Magellan, FEI) equipped with a custom-built SBEM microtome (courtesy of W. Denk(Denk and Horstmann, 2004)). Fast continuous imaging mode was implemented by adding piezo actors to operate in line with geared motors(Schmidt et al., 2017). For the acquired dataset, the sample position was centered to the C2 barrel (see above), and acquisition was started in continuous imaging mode for field of views of 229 x 217 µm^2^ each, separated by motor movements to yield a 2x2 tiling of the 453.2 x 430.4 μm^2^ region (including overlaps) imaged at 11.24 nm in-plane resolution. Each of these continuously imaged field of views was considered one “motortile” (MT1-4). Subsequently, 12,000 consecutive image planes were acquired, interleaved by microtome cuts set to a cutting thickness of 30 nm, yielding a total extent of the dataset in the cutting direction of 360 μm (Fig. 1D). The incident electron energy was 2.8 keV, beam current 3.2 nA and dwell time 100 ns, resulting in a dose of approximately 16 electrons nm^−2^. The effective data acquisition speed, including movement overheads, was 4.67 M voxel/s. Focus and astigmatism were constantly monitored and adjusted using custom-written autofocus routines.

### Image alignment

All images obtained from one image plane and a given motortile (MT, that is, 10 × 7=70 images with a size of 2048 × 3072 voxels each) were aligned separately using a custom MATLAB (Mathworks, USA) script based on detected Speeded Up Robust Features (SURF)(Bay et al., 2008) from the overlap regions of adjacent image pairs (as described in detail for the P25 dataset of ref.(Schmidt et al., 2017)). To match aligned images from consecutive planes, a region with a size of 70% of the horizontal MT size and 50% of the vertical MT size, located at the MT center, was cross-correlated with the same region from the next image plane. The translation vector between the cross-correlation peaks was applied to the second image, respectively. In a first alignment, all 12,000 sections were aligned (360 µm in cortical axis) in MT2-4. MT1 was of lower data quality and not further processed. In MT2-4, somata were annotated and dendrites reconstructed for cell type definition. For axon reconstruction and synapse identification, a higher-quality alignment was performed on the first 10,000 sections (300 µm in cortical axis). Here, debris-covered slices were manually detected and excluded from alignment, yielding a total of 9855 slices. After alignment, the 4 resulting 3-D image stacks for each of the 4 MTs of the dataset were each converted to the webKnossos data format (see https://webknossos.org) by splitting the data into data cubes with a size of 128×128×128 voxels each. These data were then uploaded to our online data annotation software webKnossos(Boergens et al., 2017) for in-browser distributed data visualization, neurite skeletonization and synapse identification.

### Volume and neuron density measurements

The volume of the acquired dataset was 453.2 x 430.4 x 360 µm^3^ (each MT was 229x217 µm, with overlaps of 4.8 µm and 3.6 µm in x and y direction, respectively). Soma density in the first 300 µm of MTs 2,3,4 was on average 110,225 per mm^3^. The final 60 µm increased soma numbers by factor 1.178 (slightly lower soma density in beginning of L5A). Together, the entire imaged volume contained 7,598 neurons. Of these, 5,259 were explicitly counted. When including glial somata, neuron fraction was found to be 69.9% and 71.9% in MT2 and MT3, respectively.

### Identification of cell types

We identified 1976 cell bodies in the fraction of dataset with optimal staining and image alignment (Fig. 1G). For the classification of ExNs and INs, we initially used the size of the cell body and rate of mitochondria in the cytosol to identify INs, which had large and mitochondria-rich cell bodies (n=127 INs). We then confirmed that these INs had low spine rates along their dendrites. To determine whether any of the small-soma neurons could also be INs, we checked their dendrites and identified another n=5 cells with low spine rates but small soma size, yielding n=132 INs total (6.7%). INs frequently had axons that exited from the cell body or from primary dendrites with an orientation towards the pia.

We then determined for all ExNs whether they had an apical dendrite by reconstructing vertically oriented primary dendrites exiting the soma towards the pia. To distinguish between L3 pyramidal and L4 star pyramidal neurons, we determined the cortex depth at which the first fluorescent cell bodies occurred as the beginning of L4 (Fig. 1G; quantified post-hoc as the 10^th^ percentile of a sigmoid fit to the cumulated fluorescent cell bodies along cortical depth based on n=160 fluorescent somata). To distinguish between L4 star pyramidal cells and L5A pyramidal neurons, we determined the depth at which no more apical-dendrite-free excitatory neurons (i.e., spiny stellates) were found as the beginning of L5A. This was confirmed post-hoc by determining the depth of steepest increase in soma density from white matter towards pia (Fig. 1G). 117 somata, positioned outside the barrel, had incomplete dendritic trees and could therefore not be classified; these were excluded from further analysis.

### Definition of thalamocortical axons

To identify the axons that were likely originating from the thalamus we used the fact that thalamic axons from VPM have been described to establish large multi-synaptic boutons at high frequency in mouse S1 cortex(Bopp et al., 2017; Motta et al., 2019) (Suppl. Fig. 4A). We applied these criteria and identified 20 such TC axons (Fig. 6), entering from the direction of the white matter.

### Synapse identification

Synapses were identified by following the trajectory of axons in webKnossos(Boergens et al., 2017). First, vesicle clouds in the axon were identified as accumulations of vesicles. Subsequently, the most likely postsynaptic target was identified by the following criteria: direct apposition with vesicle cloud; presence of a darkening and broadening of the synaptic membrane, indicative of a postsynaptic density; vesicles very close to the membrane at the site of contact. Criteria were similar to those in refs. (Karimi et al., 2020; Schmidt et al., 2017; Staffler et al., 2017). Synapses were marked as uncertain whenever the signs of darkened postsynaptic density could not be clearly identified. All analysis in this study were conducted only on synapses that had been classified as certain. The resulting axonal synapse densities were 0.23±0.50 per μm (TC axons, mean±s.d., n=20) and 0.16±0.04 per μm (IN axons, mean±s.d., n=18).

### Synapse size estimation

A random subset of synapses from TC axons to excitatory cells (n=17 for StP, n=40 for SpS) and interneurons (n=20 for W-BINs. N=20 for L-BINs, n=11 for L4 INs) were used to determine the size of the post-synaptic density (PSD). An expert annotator placed an edge along the longest dimension of the synapse which was used to estimate the PSD size for comparison across post-synaptic groups (see Suppl. Fig. 4D,E).

### Definition of Barrel volume

To evaluate axonal projections in the context of the cortical column and L4, we needed to define the outlines of the home (C2) barrel. For this, the 3D fluorescence-LM dataset comprising 600 image tiles (256 x 256 voxels each) outlining the C2 barrel was used. Image tiles 320 to 550 were used for barrel surface reconstruction using morphological operations on subsequent image planes. The results were stitched into a global reference frame followed by Gaussian smoothing (kernel 3 x 3 x 3 voxel, s.d. 2 voxels) to obtain the 3D isosurface (threshold 0.5) as 3D convex hull. Using a coarse LM-EM transformation based on correspondence points in blood vessels, an affine transformation was computed to convert the LM barrel isosurface into 3D EM reference space (3 pairs of nodes at bifurcations in blood vessels and 13 pairs of nodes in cell bodies, defined by an expert annotator (see supplementary data: tform_map_MT4.xlsx) using Horn’s quaternion-based method (Horn, 1987). Note that this definition of the barrel (and consequently its lower and upper boundary) was used for all analyses in Fig. 2-6. The definition of layer borders as a simplified tangential plane (based on fluorescent soma density along the cortical axis, as described above) was only used for Fig. 1 to obtain a conservative estimate of cell subtype distributions along the cortical axis (Fig. 1G).

### Quantitative Analysis of Axonal Projections

Based on the reconstruction of the axonal projection of L4 INs, we calculated the fraction of the axon that was confined to a barrel (“axon barrel ratio”) and the fraction that was confined to L4 (“axon L4 ratio”)(Koelbl et al., 2015). The “axon barrel ratio” of the axonal projection was calculated as the ratio between the axonal path length of the L4 IN that was within the home barrel (C2) of the neuron (defined by the barrel isosurface in EM reference space) and the total axon length. The “axon L4 ratio” of the axonal projection was calculated as the ratio between the axonal path length of the L4 IN that was within the L4 borders and the total axon length. The L4 borders (Fig. 1G) were defined as described above.

### Interneuron subtype definition

Interneurons with axonal path length less than 1.5 mm were excluded from this analysis. Hierarchical clustering (Euclidean distance, Ward’s method) was used to determine TC INs cluster (Fig. 2) as well as to separate subtypes within the TC INs (Fig. 3).

### Axon diameter measurements

TC axonal diameters (Fig. 6H) were reconstructed from three TC axons by measuring the diameters at random locations along its myelinated and unmyelinated trajectory (n=58, 39 and 32 contour annotations for the three axons, respectively). Contours were traced in the orthogonal viewport most perpendicular to the local axon axis in webKnossos. In case of myelinated segments, the inner unmyelinated axon diameter was measured and contour was labelled as “myelinated” (n=5, 7 and 5 for the three axons, red dots in Fig. 6H inset). In case of branch points the diameter was measured before the branch point. The diameter measurements were linearly interpolated to get the average distribution (bold black curve) shown in Fig 6H. The EM cross sections in Fig. 6i were chosen at distances 44 and 122 μm (arrows Fig. 6H) for one of the TC axons.

### Local circuit cFFI analysis

Local circuit analysis was based on the connectivity data reporting the number of synapses between pairs of neurons or neurites (see supplementary data extractConnectome.mat) in the dataset. The synaptic connectivity between 3 TC axons, 558 ExN (175 StP, 383 SpS) and 11 fully reconstructed INs (types whole-barrel, lower-barrel and L4-related) that had their soma in the dataset was analyzed. The connectivity matrix between these neurons was binarized to represent the presence or absence of synaptic connectivity. For each TC axon, all of its excitatory targets were inspected for an additional inhibitory connection from the targeted interneurons by the same TC axon. After excluding excitatory targets with their soma outside the barrel region, the resulting number of cFFI configurations with the additional IN-to-ExN connection were found to be 103 (41 (8 StP and 33 SpS cells), 45 (10 StP and 35 SpS cells) and 17 (5 StP and 12 SpS cells) for the 3 TC axons, respectively) and those without the IN-to-ExN connection were 5 (1 (StP cell), 1 (SpS cell) and 3 (1 StP and 2 SpS cells) for the 3 TC axons), yielding a 95% (92% StP and 96% SpS cells) cFFI ratio (see Fig. 6F). When correcting for border effects by excluding excitatory targets near the border of the dataset which limits the complete dendrite reconstruction, the cFFI was found to be 100%. Total synapses from the 3 TC axons to excitatory targets were 59, 53 and 23 and to IN targets were 4, 4 and 9, respectively. Total synapses from all the INs onto excitatory targets were 139, 179 and 50 respectively within the cFFI configurations for 3 TC axons.

### Analysis of TC synapse clustering

The axons making TC synapse onto WB-BINs and L-BINs were locally checked by an expert annotator for presence of additional synapses onto same dendritic target. This was done for all TC synapses received by W-BINs and L-BINs and used to determine fraction of clustered TC –to- IN synapses.

### IN-to-IN connectivity

To determine the IN-to-IN connectome (Fig. 4B), all output synapses of the reported INs were inspected with all postsynaptic IN dendrites present in webKnossos, such that any synapse onto any other IN was detected. Therefore, the “zero” entries in the connectome represent no synaptic connection in the dataset.

### Statistical tests

All statistical tests were performed using MATLAB and the Statistics Toolbox Release 2021a (MathWorks, USA). All tests were done using Wilcoxon-rank-sum test unless otherwise noted.

## AUTHOR CONTRIBUTIONS

MH conceived, initiated and supervised the study, YH conducted EM experiments with contributions by PL using 3D EM methodology contributed by KMB; YH, SL, MH analysed structural data; JK supervised and YH, VP conducted functional recordings and analysis. YH, SL, MH wrote the paper with contributions by all authors.

## ACKNOWLEDGEMENTS

We thank A Motta for comments on an earlier version of the manuscript, I Wolf for staining support, S Babl, L Bezzenberger, R Jakoby, R Kneißl and M Kronawitter for annotator training and task management, H Wissler for tracer management and support with figure generation, and A. Kessler, A. Brandt, A. J. Lopez, A. Strubel, A. Weyh, A. Kolbinger, A. Fantur, A. K. Spohner, A. Rix, A. Possmayer, A. Bamberg, A. Machel, A. Al-Shaboti, B. Kuhl, B. Heftrich, B. L. Stiehl, C. Lossnitzer, C. A. Sandhof, C. Arras, C. Sabatelli, C. Schumm, C. Kurz, D. Beyer, D. Baltissen, D. Greco, D. Celik, D. Kurt, D. Acay, D. J. Scheliu, E. Laube, F. Gohl, F. Sahin, F. Y. Basoeki, F. Krämer, H. Charif, H. Hees, I. Meindl, I. Stasiuk, I. Metz, J. Winkelmeier, J. Hartel, J. Meyer, J. E. Martínez, J. Knauer, J. Depnering, J. Heller, J. Kubat, J. Buss, K. Kramer, K. Strahler, K. Trares, K. Desch, K. Lust, L. Buxmann, L. Matzner, L. Kirchner, L. Praeve, L. Kreppner, L. Schütz, L. Bezzenberger, L. Hartwig, L. Lutz, L. Burchartz, M. Aly, M. Werr, M. Dell, M. Nonnenbroich, M. Harwardt, M. Mittag, M. Groothuis, M. Präve, M. Karabel, M. Weber, N. Cipta, N. Zeitzschel, N. Aydin , N. Berghaus, N. Böffinger, O. Ilea, P. König, P. Remmele, P. Müller, P. Werner, R. Gebauer, R. Hülse, R. Thieleking, S. Reichel, S. Mehlmann, S. T. Stahl, S. Umbach, S. Wehrheim, S. Bohne, S. Reibeling, S. Lerchl, T. Köcke, T. Ernst, T. Decker, T. Engelmann, T. Winkelmeier, T. Hörmann, T. Reimann, V. Kalbert, V. Pantle, V. Robl, V. Gosch, V. Dienst, W. Pfohl, Y. Wang, Yan Lu, Haibin Sheng, Yumeng Qi, and Shengxiong Wang for neuron reconstructions.

## REFERENCES

1. Agmon, A., and Connors, B.W. (1991). Thalamocortical responses of mouse somatosensory (barrel) cortexin vitro. Neuroscience 41, 365–379.

2. Ahmed, B., Anderson, J.C., Douglas, R.J., Martin, K.A.C., and Nelson, J.C. (1994). Polyneuronal innervation of spiny stellate neurons in cat visual cortex. Journal of Comparative Neurology 341, 39–49.

3. Ahmed, B., Anderson, J.C., Martin, K.A.C., and Nelson, J.C. (1997). Map of the synapses onto layer 4 basket cells of the primary visual cortex of the cat. Journal of Comparative Neurology 380, 230–242.

4. Alger, B., and Nicoll, R. (1982). Feed-forward dendritic inhibition in rat hippocampal pyramidal cells studied in vitro. The Journal of physiology 328, 105–123.

5. Arabzadeh, E., Zorzin, E., and Diamond, M.E. (2005). Neuronal Encoding of Texture in the Whisker Sensory Pathway. PLOS Biology 3, e17.

6. Ascoli, G.A., Alonso-Nanclares, L., Anderson, S.A., Barrionuevo, G., Benavides-Piccione, R., Burkhalter, A., Buzsáki, G., Cauli, B., DeFelipe, J., Fairén, A., et al. (2008). Petilla terminology: nomenclature of features of GABAergic interneurons of the cerebral cortex. Nature Reviews Neuroscience 9, 557–568.

7. Bagnall, Martha W., Hull, C., Bushong, Eric A., Ellisman, Mark H., and Scanziani, M. (2011). Multiple Clusters of Release Sites Formed by Individual Thalamic Afferents onto Cortical Interneurons Ensure Reliable Transmission. Neuron 71, 180–194.

8. Bay, H., Ess, A., Tuytelaars, T., and Van Gool, L. (2008). Speeded-Up Robust Features (SURF). Computer Vision and Image Understanding 110, 346–359.

9. Beierlein, M., Gibson, J.R., and Connors, B.W. (2000). A network of electrically coupled interneurons drives synchronized inhibition in neocortex. Nature Neuroscience 3, 904–910.

10. Beierlein, M., Gibson, J.R., and Connors, B.W. (2003). Two Dynamically Distinct Inhibitory Networks in Layer 4 of the Neocortex. Journal of Neurophysiology 90, 2987–3000.

11. Boergens, K.M., Berning, M., Bocklisch, T., Braunlein, D., Drawitsch, F., Frohnhofen, J., Herold, T., Otto, P., Rzepka, N., Werkmeister, T., et al. (2017). webKnossos: efficient online 3D data annotation for connectomics. Nat Methods 14, 691–694.

12. Bopp, R., Holler-Rickauer, S., Martin, K.A.C., and Schuhknecht, G.F.P. (2017). An Ultrastructural Study of the Thalamic Input to Layer 4 of Primary Motor and Primary Somatosensory Cortex in the Mouse. The Journal of Neuroscience 37, 2435–2448.

13. Braak, H. (1976). On the striate area of the human isocortex. A golgi- and pigmentarchitectonic study. Journal of Comparative Neurology 166, 341–364.

14. Brecht, M., and Sakmann, B. (2002). Dynamic representation of whisker deflection by synaptic potentials in spiny stellate and pyramidal cells in the barrels and septa of layer 4 rat somatosensory cortex. J Physiol 543, 49–70.

15. Brodmann, K. (1909). Vergleichende Lokalisationslehre der Grosshirnrinde in ihren Prinzipien dargestellt auf Grund des Zellenbaues (Leipzig: Johann Ambrosius Barth).

16. Bruno, R.M., and Sakmann, B. (2006). Cortex Is Driven by Weak but Synchronously Active Thalamocortical Synapses. Science 312, 1622–1627.

17. Bruno, R.M., and Simons, D.J. (2002). Feedforward Mechanisms of Excitatory and Inhibitory Cortical Receptive Fields. The Journal of Neuroscience 22, 10966–10975.

18. Buzsáki, G. (1984). Feed-forward inhibition in the hippocampal formation. Progress in neurobiology 22, 131–153.

19. Campbell, A.W., Schlesinger, E.B., and Riley, H.A. (1905). Histological studies on the localisation of cerebral function (Cambridge: University Press).

20. Carr, C., and Konishi, M. (1990). A circuit for detection of interaural time differences in the brain stem of the barn owl. The Journal of Neuroscience 10, 3227–3246.

21. Carvell, G.E., and Simons, D.J. (1990). Biometric analyses of vibrissal tactile discrimination in the rat. Journal of Neuroscience 10, 2638–2648.

22. Clark, W.E. (1925). The Visual Cortex of Primates. J Anat 59, 350–357.

23. Collins, C.E., Hendrickson, A., and Kaas, J.H. (2005). Overview of the visual system of tarsius. The Anatomical Record Part A: Discoveries in Molecular, Cellular, and Evolutionary Biology 287A, 1013-1025.

24. Connors, B.W. (2017). Synchrony and so much more: Diverse roles for electrical synapses in neural circuits. Developmental Neurobiology 77, 610–624.

25. Cruikshank, S.J., Lewis, T.J., and Connors, B.W. (2007). Synaptic basis for intense thalamocortical activation of feedforward inhibitory cells in neocortex. Nature Neuroscience 10, 462–468.

26. de Kock, C.P., Bruno, R.M., Spors, H., and Sakmann, B. (2007). Layer- and cell-type-specific suprathreshold stimulus representation in rat primary somatosensory cortex. J Physiol 581, 139–154.

27. Denk, W., and Horstmann, H. (2004). Serial block-face scanning electron microscopy to reconstruct three-dimensional tissue nanostructure. PLoS biology 2, e329.

28. Denk, W., Strickler, J.H., and Webb, W.W. (1990). Two-photon laser scanning fluorescence microscopy. Science 248, 73–76.

29. Duguid, I., Branco, T., Chadderton, P., Arlt, C., Powell, K., and Häusser, M. (2015). Control of cerebellar granule cell output by sensory-evoked Golgi cell inhibition. Proceedings of the National Academy of Sciences 112, 13099–13104.

30. Egger, V., Nevian, T., and Bruno, R.M. (2008). Subcolumnar dendritic and axonal organization of spiny stellate and star pyramid neurons within a barrel in rat somatosensory cortex. Cereb Cortex 18, 876–889.

31. Elston, G.N., Pow, D.V., and Calford, M.B. (1997). Neuronal composition and morphology in layer IV of two vibrissal barrel subfields of rat cortex. Cerebral Cortex 7, 422–431.

32. Feldmeyer, D., Egger, V., Lübke, J., and Sakmann, B. (1999). Reliable synaptic connections between pairs of excitatory layer 4 neurones within a single ‘barrel’ of developing rat somatosensory cortex. The Journal of physiology 521, 169–190.

33. Gabernet, L., Jadhav, S.P., Feldman, D.E., Carandini, M., and Scanziani, M. (2005). Somatosensory Integration Controlled by Dynamic Thalamocortical Feed-Forward Inhibition. Neuron 48, 315–327.

34. Gainey, M.A., Aman, J.W., and Feldman, D.E. (2018). Rapid Disinhibition by Adjustment of PV Intrinsic Excitability during Whisker Map Plasticity in Mouse S1. The Journal of Neuroscience 38, 4749–4761.

35. Galarreta, M., and Hestrin, S. (1999). A network of fast-spiking cells in the neocortex connected by electrical synapses. Nature 402, 72–75.

36. Garey, L.J., and Harris, G.W. (1971). A light and electron microscopic study of the visual cortex of the cat and monkey. Proceedings of the Royal Society of London Series B Biological Sciences 179, 21–40.

37. Gibson, J.R., Beierlein, M., and Connors, B.W. (1999). Two networks of electrically coupled inhibitory neurons in neocortex. Nature 402, 75–79.

38. Greenberg, D.S., and Kerr, J.N. (2009). Automated correction of fast motion artifacts for two-photon imaging of awake animals. Journal of neuroscience methods 176, 1–15.

39. Gutnisky, D.A., Yu, J., Hires, S.A., To, M.S., Bale, M.R., Svoboda, K., and Golomb, D. (2017). Mechanisms underlying a thalamocortical transformation during active tactile sensation. PLoS Comput Biol 13, e1005576.

40. Hassler, R., and Wagner, A. (1965). Experimentelle und morphologische Befunde über die vierfache corticale Projektion des visuellen Systems. In Eighth International Congress on Neurology, pp. 77–96.

41. Hatch, R.J., Mendis, G.D.C., Kaila, K., Reid, C.A., and Petrou, S. (2017). Gap Junctions Link Regular-Spiking and Fast-Spiking Interneurons in Layer 5 Somatosensory Cortex. Frontiers in Cellular Neuroscience 11.

42. Helmstaedter, M., Briggman, K.L., Turaga, S.C., Jain, V., Seung, H.S., and Denk, W. (2013). Connectomic reconstruction of the inner plexiform layer in the mouse retina. Nature 500, 168–174.

43. Hestrin, S., and Galarreta, M. (2005). Electrical synapses define networks of neocortical GABAergic neurons. Trends in neurosciences 28, 304–309.

44. Horn, B.K. (1987). Closed-form solution of absolute orientation using unit quaternions. Josa a 4, 629–642.

45. House, D.R., Elstrott, J., Koh, E., Chung, J., and Feldman, D.E. (2011). Parallel regulation of feedforward inhibition and excitation during whisker map plasticity. Neuron 72, 819–831.

46. Hua, Y., Laserstein, P., and Helmstaedter, M. (2015). Large-volume en-bloc staining for electron microscopy-based connectomics. Nat Commun 6, 7923.

47. Hubel, D.H., and Wiesel, T.N. (1962). Receptive fields, binocular interaction and functional architecture in the cat’s visual cortex. The Journal of physiology 160, 106–154.

48. Isaacson, Jeffry S., and Scanziani, M. (2011). How Inhibition Shapes Cortical Activity. Neuron 72, 231–243.

49. Isett, B.R., Feasel, S.H., Lane, M.A., and Feldman, D.E. (2018). Slip-Based Coding of Local Shape and Texture in Mouse S1. Neuron 97, 418–433.e415.

50. Jadhav, S.P., Wolfe, J., and Feldman, D.E. (2009). Sparse temporal coding of elementary tactile features during active whisker sensation. Nature Neuroscience 12, 792–800.

51. Kaas, J.H., Hall, W.C., and Diamond, I.T. (1972). Visual cortex of the grey squirrel (Sciurus carolinensis): Architectonic subdivisions and connections from the visual thalamus. Journal of Comparative Neurology 145, 273–305.

52. Kanichay, R.T., and Silver, R.A. (2008). Synaptic and cellular properties of the feedforward inhibitory circuit within the input layer of the cerebellar cortex. Journal of Neuroscience 28, 8955–8967.

53. Kanold, P.O., Kara, P., Reid, R.C., and Shatz, C.J. (2003). Role of Subplate Neurons in Functional Maturation of Visual Cortical Columns. Science 301, 521–525.

54. Kara, P., Pezaris, J.S., Yurgenson, S., and Reid, R.C. (2002). The spatial receptive field of thalamic inputs to single cortical simple cells revealed by the interaction of visual and electrical stimulation. Proceedings of the National Academy of Sciences 99, 16261–16266.

55. Karimi, A., Odenthal, J., Drawitsch, F., Boergens, K., and Helmstaedter, M. (2020). Cell-type specific innervation of cortical pyramidal cells at their apical dendrites. Elife 9.

56. Karnani, M.M., Jackson, J., Ayzenshtat, I., Hamzehei Sichani, A., Manoocheri, K., Kim, S., and Yuste, R. (2016). Opening Holes in the Blanket of Inhibition: Localized Lateral Disinhibition by VIP Interneurons. The Journal of Neuroscience 36, 3471–3480.

57. Koelbl, C., Helmstaedter, M., Lubke, J., and Feldmeyer, D. (2015). A barrel-related interneuron in layer 4 of rat somatosensory cortex with a high intrabarrel connectivity. Cereb Cortex 25, 713–725.

58. Kornfeld, J., Benezra, S.E., Narayanan, R.T., Svara, F., Egger, R., Oberlaender, M., Denk, W., and Long, M.A. (2017). EM connectomics reveals axonal target variation in a sequence-generating network. Elife 6, e24364.

59. Lee, S., Kruglikov, I., Huang, Z.J., Fishell, G., and Rudy, B. (2013). A disinhibitory circuit mediates motor integration in the somatosensory cortex. Nature Neuroscience 16, 1662–1670.

60. LeVay, S. (1973). Synaptic patterns in the visual cortex of the cat and monkey. Electron microscopy of Golgi Preparations. Journal of Comparative Neurology 150, 53–85.

61. Lewis, W.B., and Ferrier, D. (1880). III. Researches on the comparative structure of the cortex cerebri. Philosophical Transactions of the Royal Society of London 171, 35–64.

62. Lorente De No, R. (1938). Cerebral cortex : architecture, intracortical connections, motor projections. Physiology of the Nervous System, 288-313.

63. Lübke, J., Egger, V., Sakmann, B., and Feldmeyer, D. (2000). Columnar Organization of Dendrites and Axons of Single and Synaptically Coupled Excitatory Spiny Neurons in Layer 4 of the Rat Barrel Cortex. The Journal of Neuroscience 20, 5300–5311.

64. Lübke, J., Roth, A., Feldmeyer, D., and Sakmann, B. (2003). Morphometric Analysis of the Columnar Innervation Domain of Neurons Connecting Layer 4 and Layer 2/3 of Juvenile Rat Barrel Cortex. Cerebral Cortex 13, 1051–1063.

65. Lund, J.S. (1973). Organization of neurons in the visual cortex, area 17, of the monkey (Macaca mulatta). Journal of Comparative Neurology 147, 455–495.

66. Martinez, L.M., Wang, Q., Reid, R.C., Pillai, C., Alonso, J.-M., Sommer, F.T., and Hirsch, J.A. (2005). Receptive field structure varies with layer in the primary visual cortex. Nature Neuroscience 8, 372–379.

67. McCormick, D.A., Connors, B.W., Lighthall, J.W., and Prince, D.A. (1985). Comparative electrophysiology of pyramidal and sparsely spiny stellate neurons of the neocortex. J Neurophysiol 54, 782–806.

68. Mittmann, W., Koch, U., and Häusser, M. (2005). Feed-forward inhibition shapes the spike output of cerebellar Purkinje cells. The Journal of physiology 563, 369–378.

69. Motta, A., Berning, M., Boergens, K.M., Staffler, B., Beining, M., Loomba, S., Hennig, P., Wissler, H., and Helmstaedter, M. (2019). Dense connectomic reconstruction in layer 4 of the somatosensory cortex. Science 366.

70. Oberlaender, M., Ramirez, A., and Bruno, R.M. (2012). Sensory experience restructures thalamocortical axons during adulthood. Neuron 74, 648–655.

71. Otsuka, R., and Hassler, R. (1962). Über Aufbau und Gliederung der corticalen Sehsphäre bei der Katze. Archiv für Psychiatrie und Nervenkrankheiten 203, 212–234.

72. Park, J., Rodgers, C., Hong, Y.K., Dahan, J., and Bruno, R. (2019). Primary somatosensory cortex is essential for texture discrimination but not object detection in mice. IBRO Reports 6, S550.

73. Pernelle, G., Nicola, W., and Clopath, C. (2018). Gap junction plasticity as a mechanism to regulate network-wide oscillations. PLOS Computational Biology 14, e1006025.

74. Pfeffer, C.K., Xue, M., He, M., Huang, Z.J., and Scanziani, M. (2013). Inhibition of inhibition in visual cortex: the logic of connections between molecularly distinct interneurons. Nature Neuroscience 16, 1068–1076.

75. Porter, J.T., Johnson, C.K., and Agmon, A. (2001). Diverse Types of Interneurons Generate Thalamus-Evoked Feedforward Inhibition in the Mouse Barrel Cortex. The Journal of Neuroscience 21, 2699–2710.

76. Pouille, F., and Scanziani, M. (2001). Enforcement of temporal fidelity in pyramidal cells by somatic feed-forward inhibition. Science 293, 1159–1163.

77. Ramachandra, V., Pawlak, V., Wallace, D.J., and Kerr, J.N.D. (2020). Impact of visual callosal pathway is dependent upon ipsilateral thalamus. Nat Commun 11, 1889.

78. Ramón y Cajal, S. (1899). Estudios sobre la corteza cerebral humana. Corteza visual. Rev Trim Microgr 4, 1–63.

79. Rikhye, R.V., Yildirim, M., Hu, M., Breton-Provencher, V., and Sur, M. (2021). Reliable sensory processing in mouse visual cortex through cooperative interactions between somatostatin and parvalbumin interneurons. The Journal of Neuroscience.

80. Rudy, B., Fishell, G., Lee, S., and Hjerling-Leffler, J. (2011). Three groups of interneurons account for nearly 100% of neocortical GABAergic neurons. Developmental Neurobiology 71, 45–61.

81. Scala, F., Kobak, D., Shan, S., Bernaerts, Y., Laturnus, S., Cadwell, C.R., Hartmanis, L., Froudarakis, E., Castro, J.R., Tan, Z.H., et al. (2019). Layer 4 of mouse neocortex differs in cell types and circuit organization between sensory areas. Nature Communications 10, 4174.

82. Schmidt, H., Gour, A., Straehle, J., Boergens, K.M., Brecht, M., and Helmstaedter, M. (2017). Axonal synapse sorting in medial entorhinal cortex. Nature 549, 469–475.

83. Schubert, D., Kötter, R., Zilles, K., Luhmann, H.J., and Staiger, J.F. (2003). Cell Type-Specific Circuits of Cortical Layer IV Spiny Neurons. The Journal of Neuroscience 23, 2961–2970.

84. Simons, D.J., and Woolsey, T.A. (1984a). Morphology of Golgi-Cox-impregnated barrel neurons in rat SmI cortex. The Journal of comparative neurology 230, 119–132.

85. Simons, D.J., and Woolsey, T.A. (1984b). Morphology of Golgi-Cox-impregnated barrel neurons in rat SmI cortex. Journal of Comparative Neurology 230, 119–132.

86. Staffler, B., Berning, M., Boergens, K.M., Gour, A., Smagt, P.v.d., and Helmstaedter, M. (2017). SynEM, automated synapse detection for connectomics. Elife 6, e26414.

87. Staiger, J.F., Flagmeyer, I., Schubert, D., Zilles, K., Kotter, R., and Luhmann, H.J. (2004). Functional diversity of layer IV spiny neurons in rat somatosensory cortex: quantitative morphology of electrophysiologically characterized and biocytin labeled cells. Cereb Cortex 14, 690–701.

88. Staiger, J.F., Zuschratter, W., Luhmann, H.J., and Schubert, D. (2009). Local circuits targeting parvalbumin-containing interneurons in layer IV of rat barrel cortex. Brain Structure and Function 214, 1.

89. Stratford, K.J., Tarczy-Hornoch, K., Martin, K.A.C., Bannister, N.J., and Jack, J.J.B. (1996). Excitatory synaptic inputs to spiny stellate cells in cat visual cortex. Nature 382, 258–261.

90. Sun, Q.-Q., Huguenard, J.R., and Prince, D.A. (2006). Barrel Cortex Microcircuits: Thalamocortical Feedforward Inhibition in Spiny Stellate Cells Is Mediated by a Small Number of Fast-Spiking Interneurons. The Journal of Neuroscience 26, 1219–1230.

91. Taniguchi, H., He, M., Wu, P., Kim, S., Paik, R., Sugino, K., Kvitsani, D., Fu, Y., Lu, J., Lin, Y., et al. (2011). A Resource of Cre Driver Lines for Genetic Targeting of GABAergic Neurons in Cerebral Cortex. Neuron 71, 995–1013.

92. Temereanca, S., and Simons, D.J. (2004). Functional Topography of Corticothalamic Feedback Enhances Thalamic Spatial Response Tuning in the Somatosensory Whisker/Barrel System. Neuron 41, 639–651.

93. Torii, M., and Levitt, P. (2005). Dissociation of Corticothalamic and Thalamocortical Axon Targeting by an EphA7-Mediated Mechanism. Neuron 48, 563–575.

94. Tremblay, R., Lee, S., and Rudy, B. (2016). GABAergic Interneurons in the Neocortex: From Cellular Properties to Circuits. Neuron 91, 260–292.

95. Valverde, F. (1977). Lamination of the striate cortex. Journal of Neurocytology 6, 483–484.

96. von Bonin, G. (1942). The striate area of primates. Journal of Comparative Neurology 77, 405–429.

97. White, E.L. (1978). Identified neurons in mouse smi cortex which are postsynaptic to thalamocortical axon terminals: A combined golgi-electron microscopic and degeneration study. Journal of Comparative Neurology 181, 627–661.

98. Williams, L.E., and Holtmaat, A. (2019). Higher-Order Thalamocortical Inputs Gate Synaptic Long-Term Potentiation via Disinhibition. Neuron 101, 91–102.e104.

99. Wolfe, J., Hill, D.N., Pahlavan, S., Drew, P.J., Kleinfeld, D., and Feldman, D.E. (2008). Texture Coding in the Rat Whisker System: Slip-Stick Versus Differential Resonance. PLOS Biology 6, e215.

100. Wong, P., and Kaas, J.H. (2008). Architectonic Subdivisions of Neocortex in the Gray Squirrel (Sciurus carolinensis). The Anatomical Record 291, 1301–1333.

101. Wong, P., and Kaas, J.H. (2009). Architectonic Subdivisions of Neocortex in the Tree Shrew (Tupaia belangeri). The Anatomical Record 292, 994–1027.

102. Wong, P., and Kaas, J.H. (2010). Architectonic Subdivisions of Neocortex in the Galago (Otolemur garnetti). The Anatomical Record 293, 1033–1069.

103. Woolsey, T.A., Dierker, M.L., and Wann, D.F. (1975). Mouse SmI cortex: qualitative and quantitative classification of golgi-impregnated barrel neurons. Proc Natl Acad Sci U S A 72, 2165–2169.

104. Woolsey, T.A., and Van der Loos, H. (1970). The structural organization of layer IV in the somatosensory region (SI) of mouse cerebral cortex. The description of a cortical field composed of discrete cytoarchitectonic units. Brain Res 17, 205–242.

105. Wu, H.-P.P., Ioffe, J.C., Iverson, M.M., Boon, J.M., and Dyck, R.H. (2013). Novel, whisker-dependent texture discrimination task for mice. Behavioural brain research 237, 238–242.

106. Xu, H., Jeong, H.-Y., Tremblay, R., and Rudy, B. (2013). Neocortical Somatostatin-Expressing GABAergic Interneurons Disinhibit the Thalamorecipient Layer 4. Neuron 77, 155–167.

107. Yu, J., Gutnisky, D.A., Hires, S.A., and Svoboda, K. (2016). Layer 4 fast-spiking interneurons filter thalamocortical signals during active somatosensation. Nature Neuroscience 19, 1647–1657.

108. Yu, J., Hu, H., Agmon, A., and Svoboda, K. (2019). Recruitment of GABAergic Interneurons in the Barrel Cortex during Active Tactile Behavior. Neuron 104, 412–427.e414.

109. Yuste, R., Hawrylycz, M., Aalling, N., Aguilar-Valles, A., Arendt, D., Armañanzas, R., Ascoli, G.A., Bielza, C., Bokharaie, V., Bergmann, T.B., et al. (2020). A community-based transcriptomics classification and nomenclature of neocortical cell types. Nature Neuroscience 23, 1456–1468.

110. Zhou, X., Mansori, I., Fischer, T., Witte, M., and Staiger, J.F. (2020). Characterizing the morphology of somatostatin-expressing interneurons and their synaptic innervation pattern in the barrel cortex of the GFP-expressing inhibitory neurons mouse. Journal of Comparative Neurology 528, 244–260.

