## Supplementary Material for "Connectomic analysis of thalamus-driven disinhibition in cortical layer 4"


### Hua Loomba et al

### SUPPLEMENTARY FIGURES

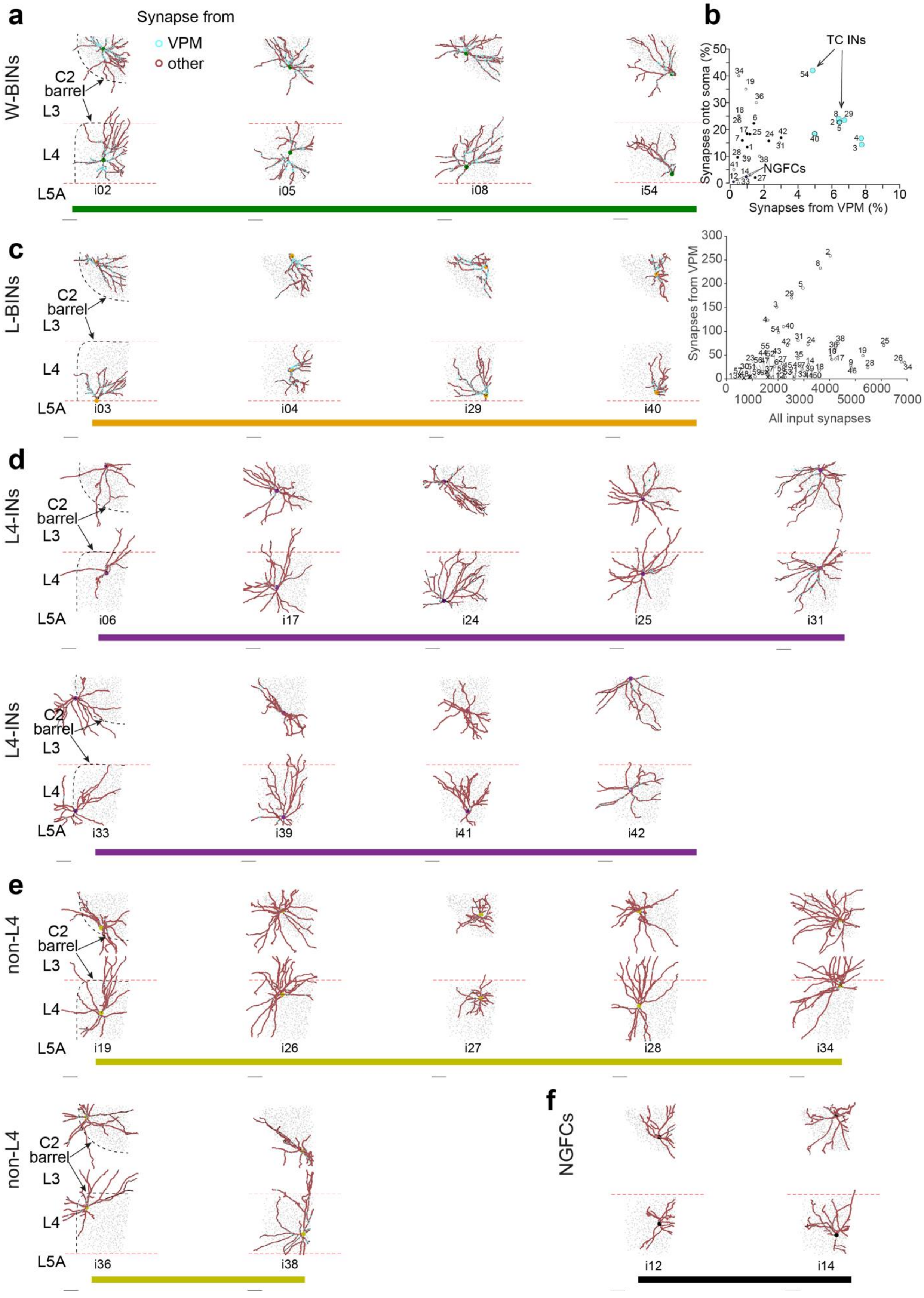

a
Synapse from
VPM
other
W-BINs
C2
barrel
L3
L4
L5A
i02
i05
i08
i54
b
TC INs
NGFCs
Synapses onto soma (%)
Synapses from VPM (%)
0
2
4
6
8
10
10
20
30
40
50
Synapses from VPM
All input synapses
0
1000
2000
3000
4000
5000
6000
7000
50
100
150
200
250
300
c
L-BINs
i03
i04
i29
i40
d
L4-INs
i06
i17
i24
i25
i31
i33
i39
i41
i42
e
non-L4
i19
i26
i27
i28
i34
i36
i38
f
NGFCs
i12
i14

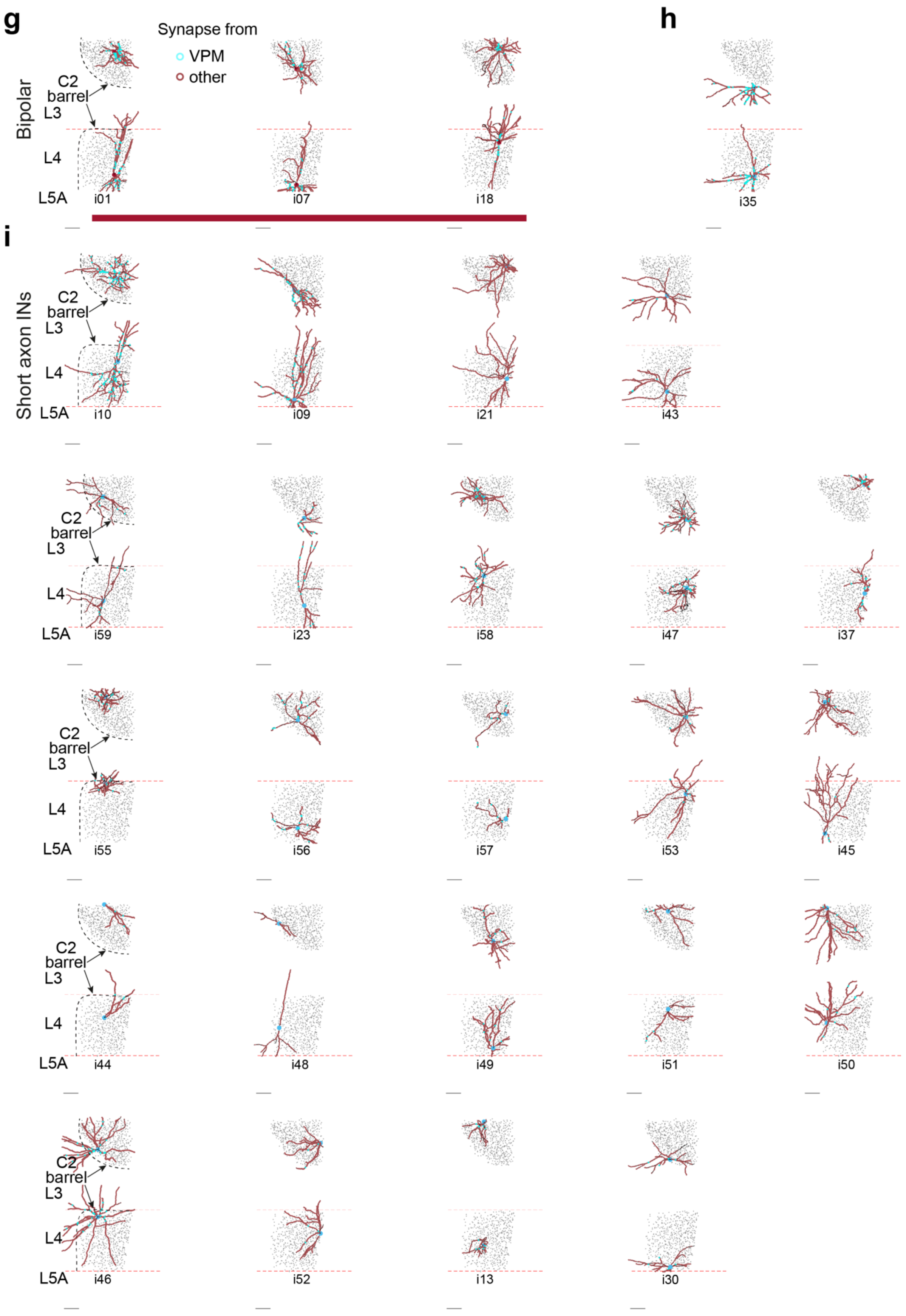
g
Synapse from
VPM
other
Bipolar
C2
barrel
L3
L4
L5A
i01
i07
i18
h
i35
i
Short axon INs
i10
i09
i21
i43
i59
i23
i58
i47
i37
i55
i56
i57
i53
i45
i44
i48
i49
i51
i50
i46
i52
i13
i30

**Supplementary Figure 1. Gallery of dendritic reconstructions of all INs with all input synapses.** IN classification as in Suppl. Fig. 2 and Suppl. Table 1. Dendrites, black; synapses from VPM, cyan, from other sources, brown. (B**)** Distribution of somatic targeting and TC input fraction over all INs (top, numbers correspond to IN index in Suppl. Fig. 1,2). Number of input synapses from VPM vs. all input synapses per IN (bottom). Crosses, INs with obviously cut dendritic tree.

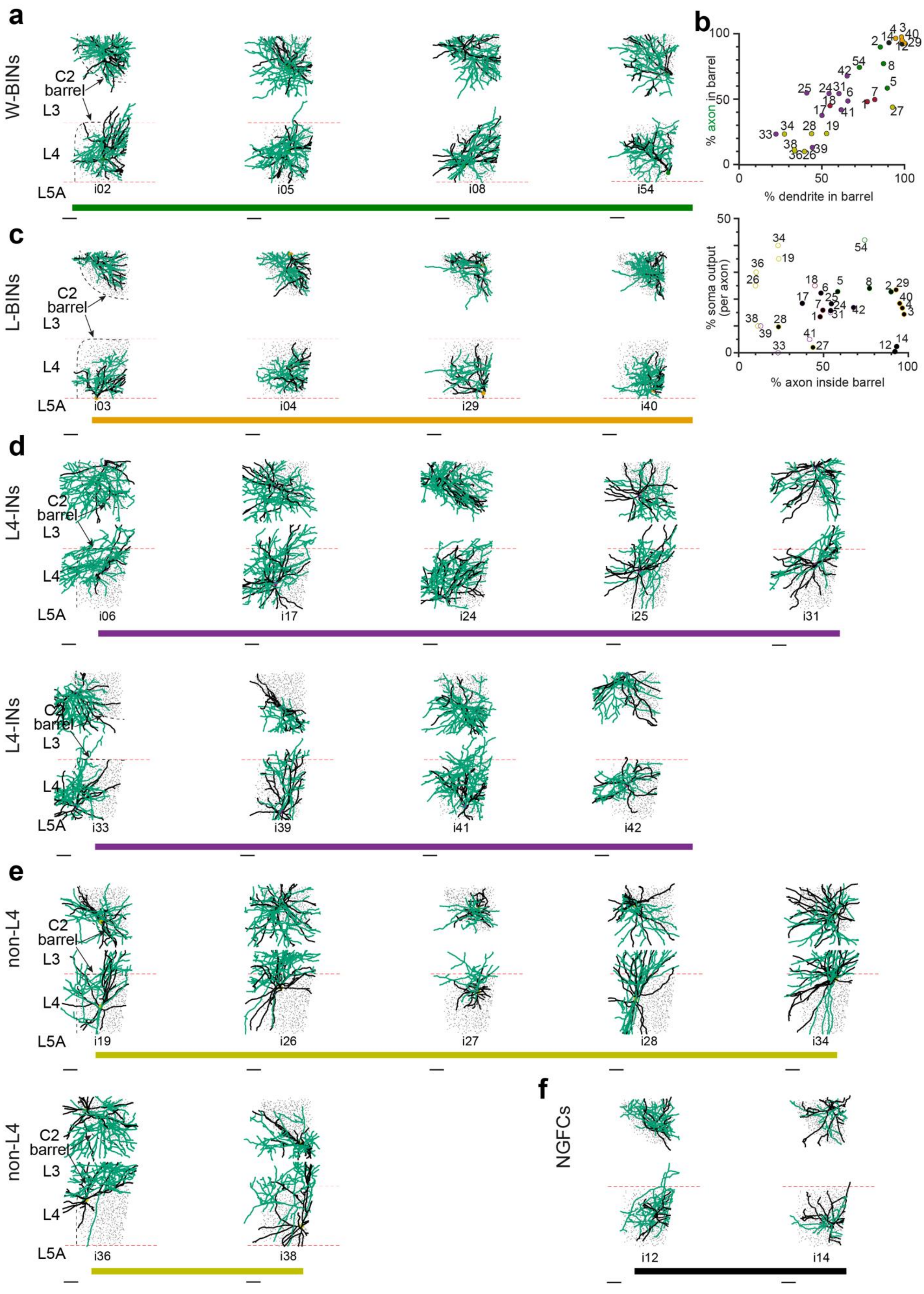

a
W-BINs
C2
barrel
L3
L4
L5A
i02
i05
i08
i54
b
% axon in barrel
% dendrite in barrel
% soma output
(per axon)
% axon inside barrel
c
L-BINs
i03
i04
i29
i40
d
L4-INs
i06
i17
i24
i25
i31
i33
i39
i41
i42
e
non-L4
i19
i26
i27
i28
i34
i36
i38
f
NGFCs
i12
i14

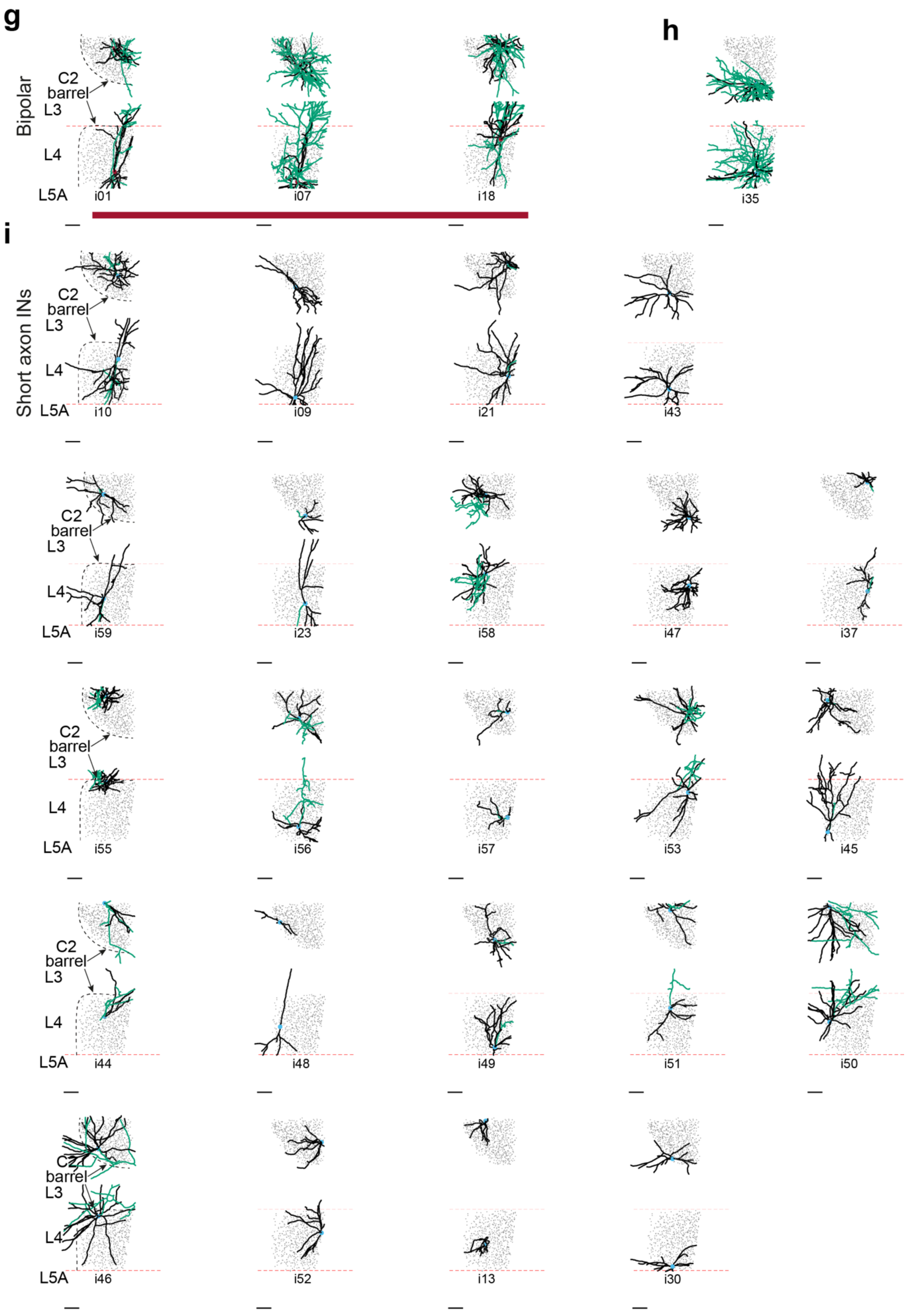
g
h
i
Bipolar
Short axon INs
C2
barrel
L3
L4
L5A
i01
i07
i18
i35
i10
i09
i21
i43
i59
i23
i58
i47
i37
i55
i56
i57
i53
i45
i44
i48
i49
i51
i50
i46
i52
i13
i30

**Supplementary Figure 2. Gallery of dendritic and axonal reconstructions of all INs.** IN classification into TC-INs (L-BIN, **a** and W-BIN, **c**) as in Fig. 2. Non-TC INs were subdivided by the projection of their axons to L4 (L4-INs, D) vs. L3 (non-L4 INs, E) for multipolar interneurons. INs with bipolar dendrites were reported as separate class (G). IN from neighboring barrel, h. INs with axon shorter than 1.5mm path length were not further classified (H). 2 INs with short local axon, no TC input and no soma output were identified as putative NGFCs (F). See Suppl. Table 1 for comparison to published IN types.

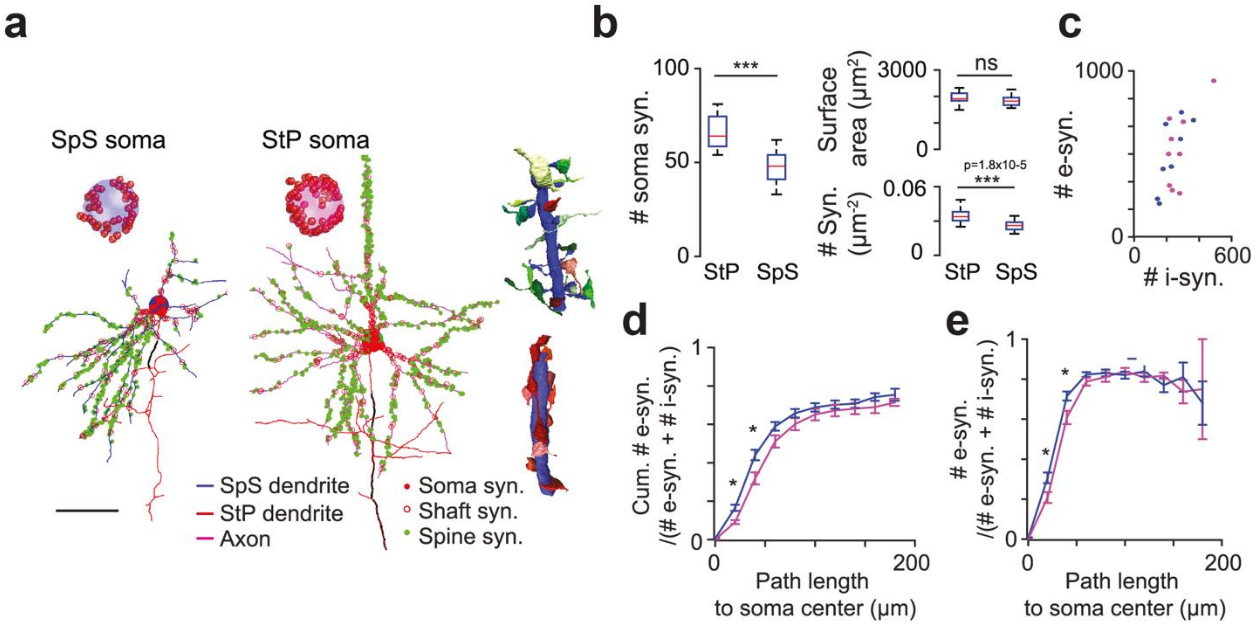


**Supplementary Figure 3. Input synapses of excitatory neurons**. **(A)** Complete mapping of somatic (red spheres), shaft (open red circles) and spine (green spheres) synapses for a SpS cell **(B)** Same as in (C) for a StP cell **(C)** Comparison of number of soma innervating synapses (StP: 66.05 ± 9.04, n = 21; SpS: 47.00 ± 8.63, n= 18, two-sample t test: $p < 0.0001$), somatic surface area (StP: 1948.2 ± 220.2 $\mu m^2$; SpS: 1828.0 ± 178.2 $\mu m^2$, two-sample t test: $p = 0.072$) and density of soma innervating synapses for StP and SpS cells (StP: 0.0343 ± 0.0061 per $\mu m^2$; SpS: 0.0258 ± 0.0045 per $\mu m^2$, two-sample t test: $p < 0.0001$). **(D)** Inhibitory soma and dendritic shafts vs. excitatory spine synapses on StP (magenta) and SpS cells (blue). **(E)** Cumulative fraction of spines of total input synapses along path length from soma centre. SpS featured a higher exc./(exc.+ihn.) ratio close to the soma than StP (for the first 20 µm, StP: 0.0913 ± 0.0302, n = 9 vs. SpS: 0.1655 ± 0.0452, n = 8, two-sample t test, $p = 0.0011$ and for the first 40 µm, StP: 0.3186 ± 0.0939 vs. SpS: 0.4404 ± 0.0770, $p = 0.0110$). **(F)** Local fraction of spines. Spines contribute about 80% (StP: 0.8130 ± 0.0189 vs. SpS: 0.8169 ± 0.0270) of all dendritic inputs at distal dendrites (>60 µm dendritic path length from soma).

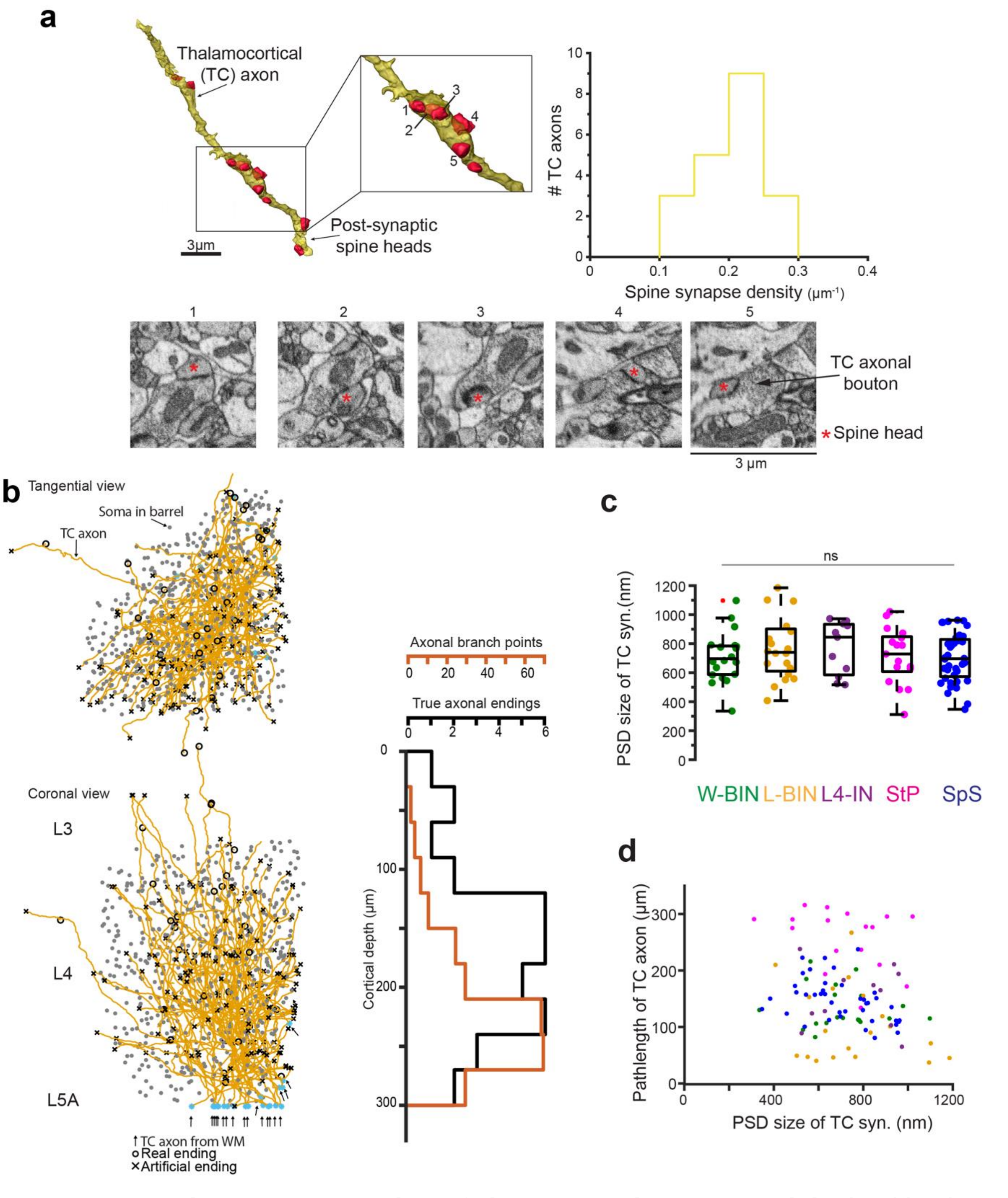


**Supplementary Figure 4. Properties of thalamocortical axons. (A)** Identification of thalamocortical axons from thalamic nucleus VPM by previously established morphological criteria in S1 mouse cortex (Bopp et al., 2017; Motta et al., 2019). Example of a TC axonal branch (top) with indication of postsynaptic spine heads. Note innervation of multiple spine heads by single axonal boutons, example of 5 spine heads targeted by one bouton shown. Right, Quantification of spine synapse density for the expert-annotated TC axons (n=20). Bottom, EM images of 5 spine heads targeted by one axonal bouton (see d). Red asterisks, postsynaptic spine heads of same TC bouton (scale bar 3 µm). **(B)** Distribution of axonal endings (black) and branchpoints (red) over cortex depth for n=20 TC axons (left). **(C)** Comparison of postsynaptic density (PSD) size (measured as largest diameter of synaptic area) of TC synapses onto W-BINs (green), L-BINs (orange), L4-INs (purple), StP (magenta) and SpS (blue) cells. **(D)** Distribution of TC synapse sizes along the axonal path length versus PSD sizes. No significant difference in synapse sizes was found (pairwise Wilcoxon rank-sum test, $p>0.05$ for each post-synaptic pair).

**Supplementary Table 1. Comparison of IN types to classification in previous studies.**

| This study | | Potential correspondence to IN types from previous studies | | | | |
|---|---|---|---|---|---|---|
| **IN IDs (see Suppl. Figs. 1,2)** | **IN Types** | **Scala et al. 2020** | **Yu et al. 2019** | **Feldmeyer et al. 2018** | **Koelbl et al. 2015** | **Tremblay et al. 2016** |
| 2 | TC-IN (W-BIN) | LBC | FS | L4 BC | Cluster 3 (BINs) | PV |
| 5 | TC-IN (W-BIN) | LBC | FS | L4 BC | Cluster 3 (BINs) | PV |
| 8 | TC-IN (W-BIN) | LBC | FS | L4 BC | Cluster 3 (BINs) | PV |
| 54 | TC-IN (W-BIN) | LBC | FS | L4 BC | Cluster 3 (BINs) | PV |
| 3 | TC-IN (L-BIN) | LBC | FS | L4 BC | Cluster 3 (BINs) | PV |
| 4 | TC-IN (L-BIN) | LBC | FS | L4 BC | Cluster 3 (BINs) | PV |
| 29 | TC-IN (L-BIN) | LBC | | L4 BC | Cluster 3 (BINs) | PV |
| 40 | TC-IN (L-BIN) | LBC | FS | L4 BC | Cluster 3 (BINs) | PV |
| 6 | L4 IN | NMC | | L4 nFS | Cluster 2 | NMC |
| 17 | L4 IN | NMC | | L4 nFS | Cluster 2 | NMC |
| 24 | L4 IN | NMC | | L4 nFS | Cluster 2 | NMC |
| 25 | L4 IN | NMC | | L4 nFS | Cluster 2 | NMC |
| 31 | L4 IN | NMC | | L4 nFS | Cluster 2 | NMC |
| 33 | L4 IN | NMC | SST | | Cluster 2 | NMC |
| 39 | L4 IN | NMC | SST | | Cluster 2 | NMC |
| 41 | L4 IN | NMC | SST | L4 nFS | Cluster 2 | NMC |
| 42 | L4 IN | NMC | | L4 nFS | Cluster 2 | NMC |
| 12 | NGFC | NGFC | | L4 NGFC | Cluster 3 (BINs) | NGFC |
| 14 | NGFC | NGFC | | L4 NGFC | Cluster 3 (BINs) | NGFC |
| 1 | BP | | | VIP | **?** | VIP |
| 7 | BP | | | VIP | **?** | VIP |
| 18 | BP | | | VIP | **?** | VIP |
| 19 | non-L4 | | | | Cluster 1 | |
| 26 | non-L4 | | | | Cluster 1 | |
| 27 | non-L4 | | | | Cluster 1 | |
| 28 | non-L4 | | | | Cluster 1 | |
| 34 | non-L4 | | | | Cluster 1 | |
| 36 | non-L4 | | | | Cluster 1 | |
| 38 | non-L4 | | | | Cluster 1 | |
| 11, 15, 16, 20, 22, 35 | neighboring barrel | | | | | |
| 9, 10, 13, 21, 23, 30, 37, 43-51 | Not classified, axon <1.5mm | | | | | |

List of all INs in L4, this study, with assigned classes and potential correspondence to previously employed classifications. LBC, large basket cell; NMC, non-Martinotti cell; NGFC, neurogliaform cell; FS, fast spiking IN; BC, basket cell; nFS, non-fast spiking IN. Comparisons refer to references (Feldmeyer et al., 2018; Koelbl et al., 2015; Scala et al., 2019; Tremblay et al., 2016; Yu et al., 2019). Total counts: n=52 INs within C2 barrel (within a volume corresponding approximately to 1/4$^{th}$ of entire C2 barrel, see Fig. 1);

n=23 INs with axon <1.5 mm path length; n=6 INs outside of C2; n=58 total identified IN somata.

Bopp, R., Holler-Rickauer, S., Martin, K.A., and Schuhknecht, G.F. (2017). An Ultrastructural Study of the Thalamic Input to Layer 4 of Primary Motor and Primary Somatosensory Cortex in the Mouse. J Neurosci 37, 2435-2448.
Feldmeyer, D., Qi, G., Emmenegger, V., and Staiger, J.F. (2018). Inhibitory Interneurons and their Circuit Motifs in the Many Layers of the Barrel Cortex. Neuroscience 368, 132-151.
Koelbl, C., Helmstaedter, M., Lubke, J., and Feldmeyer, D. (2015). A barrel-related interneuron in layer 4 of rat somatosensory cortex with a high intrabarrel connectivity. Cereb Cortex 25, 713-725.
Motta, A., Berning, M., Boergens, K.M., Staffler, B., Beining, M., Loomba, S., Hennig, P., Wissler, H., and Helmstaedter, M. (2019). Dense connectomic reconstruction in layer 4 of the somatosensory cortex. Science 366.
Scala, F., Kobak, D., Shan, S., Bernaerts, Y., Laturnus, S., Cadwell, C.R., Hartmanis, L., Froudarakis, E., Castro, J.R., Tan, Z.H., *et al.* (2019). Layer 4 of mouse neocortex differs in cell types and circuit organization between sensory areas. Nature Communications 10, 4174.
Tremblay, R., Lee, S., and Rudy, B. (2016). GABAergic Interneurons in the Neocortex: From Cellular Properties to Circuits. Neuron 91, 260-292.
Yu, J., Hu, H., Agmon, A., and Svoboda, K. (2019). Recruitment of GABAergic Interneurons in the Barrel Cortex during Active Tactile Behavior. Neuron 104, 412-427.e414.
